# Managing the genetic diversity of *Dioscorea* yams for adaptation to climate change

**DOI:** 10.64898/2026.09.10.750615

**Authors:** Bindu Adhikari, Roland Akakpo, Anna Halpin-McCormick, Marybel Soto Gomez, Nora Scarcelli, Karine Alix, Yves Vigouroux, Michael Kantar

## Abstract

Understanding adaptation to the environment is fundamental to agriculture. We analyzed a georeferenced collection of 167 accessions of three *Dioscorea* species (cultivated *D. rotundata* and its wild relatives *D. praehensilis* and *D. abyssinica*) spanning a wide range of environments, from dry Sahelian to rainforest. After accounting for species identity, population structure and geography, environment explained 1.8% of genetic variation, and genome–environment association analyses identified few loci (16 in the full panel and three within cultivated *D. rotundata*, including one soil-pH marker detected in both). Bayesian sparse linear mixed models attributed variation in annual and dry-quarter precipitation and in soil pH to a modest number of loci of larger effect. This pattern is consistent with a mixed architecture, with a few larger-effect loci on a polygenic background, which could support multiple breeding strategies. Environmental genomic prediction was accurate under random cross-validation (r ≈ 0.7) but largely reflected population structure and was not accurate for accessions from geographic areas excluded from the training data. Moreover, a core collection selected from genotypes performed as well as random training sets of the same size thus candidate parents identified here therefore require field validation. Genomic offsets under a high-emission climate scenario (SSP5-8.5) were highest for the Cameroonian lineage of *D. praehensilis*, but projected climates lie largely outside the range of climates sampled. Existing genebank collections of wild and cultivated yams contain environment-associated variation that could be used through population improvement and targeted introgression, once candidate loci and parents have been validated in the field.

**Plain Language Summary:** Yam is an ancient staple crop grown across the tropics, yet much remains unknown about how yam species adapt to their environments. We studied 167 wild and cultivated yam samples from West Africa, spanning dry Sahelian to rainforest conditions, and explored the relationship between their genomes and local climate and soil. After separating the effects of species differences and geography, the environment explained only a small part of the genetic differences among samples, and adaptation to rainfall and soil acidity appears to involve a modest number of genes. We showed that a small “core” set of samples representing all genetic groups can stand in for the full collection when predicting adaptation, and we flagged candidate parents for breeding stress-tolerant yam that now need field testing. Together, these findings give breeders starting points for developing climate-resilient yam varieties from existing genebank collections.

## Introduction

The genus *Dioscorea* comprises more than 600 species distributed across temperate, tropical and subtropical regions, and contains major and minor food crops (Ayensu and Coursey, 1972; Wilkin et al., 2005; Royal Botanic Gardens, Kew, n.d.). These species are valued for their starchy tubers or “yams” (Obidiegwu et al., 2020) and were domesticated independently in multiple regions across the world. These regions include West Africa for *Dioscorea rotundata* (white yam; 2n =2x =40) and *D. cayenensis* (yellow yam; 2n = 2x = 40, 2n =3x = 60, and 2n = 4x = 80), likely Southeast Asia for *D. alata* (water yam; 2n = 2x = 40, 2n =3x = 60, and 2n = 4x = 80), and parts of Central and South America for *D. trifida* (2n = 2x = 40, and 2n = 4x = 80) (Dansi et al., 2000; Scarcelli et al., 2019a; Sugihara et al., 2020; Sugihara et al., 2021; da Silva Souza et al., 2024; Wang et al., 2024; Zoundjihekpon et al., 1990; Obidiegwu et al., 2009). These domestication events occurred more than 5,000 years ago, making yam an ancient, cultivated staple crop (Obidiegwu & Akpabio, 2017). In West Africa, yam cultivation began 4000-5000 years ago (Shaw, 1972), with evidence of domestication in the Niger River basin region (Scarcelli et al., 2019a). The common species are semi-perennial or perennial but are typically cultivated as annuals (Rubatzky and Yamaguchi, 1997; Mignouna et al., 2009). Yam breeding has traditionally focused on clonal propagation methods(Asiedu et al., 2003). Recently, molecular breeding techniques have been increasingly used to improve the efficiency of developing yam cultivars with higher yield, disease resistance, and stress tolerance, driven by an increased number of genomic resources and genetic mapping studies (Norman et al., 2023; Adjei et al., 2024).

Cultivated yam species have wide geographical distributions (from coastal regions to savannah to forest) and many different ecological adaptations (e.g. a 1100mm rainfall gradient) that provide an opportunity to find genetic diversity (Neina, 2021). West Africa is the primary center of yam production, contributing over 90% production of *D. rotundata* and *D. cayenensis* (Darkwa et al., 2020). *Dioscorea alata* is the most widely cultivated yam species, grown across Asia, Oceania, the Caribbean, and parts of Africa due to its adaptability, high yield potential and long postharvest life (Lebot et al. 2023). *Dioscorea trifida* remains a local but key staple in the Amazonian basin of South America because of its resilience to humid tropical environments (Ngo Ngwe et al., 2015; Scarcelli et al., 2017). Yams tend to thrive in humid and sub-humid lowlands, with optimal cultivation in tropical and subtropical climates characterized by well-distributed rainfall and temperatures ranging from 25°C to 30°C (Aighewi et al., 2022). *Dioscorea rotundata* and *D. cayenensis* prefer fertile, well-drained soils with adequate moisture, while *D. alata* is adaptable to a range of environmental conditions, including varying photoperiods and temperatures (Marcos et al., 2009). Environmental change, specifically increases in temperature and soil degradation, is a major threat to production of *D. rotundata* in west Africa (Kouakou et al., 2025). The diversity of *Dioscorea* species and of the environments in which they grow is a potentially useful source of climate adaptation, but it has not been investigated systematically.

In major production regions *Dioscorea* species co-occur and exhibit interspecific compatibility (Mondo et al., 2022; Oyedoyin et al., 2025). Plant breeding efforts have drawn on this compatibility through interspecific hybridization and global germplasm collections (Darkwa et al., 2020; Mondo et al., 2022; Messadia et al., 2026). Despite this history, limited efforts have been made to breed specifically for climate resilience, a gap that is compounded by the fact that the environmental niches and underlying genetic diversity remain unknown for most yam species (except for *D. cayenensis* and *D. alata*; Pironon et al., 2019), making it difficult to operationalize the widespread diversity available in collections. One approach to operationalize collections is genome-environment association (GEA) analysis, a wide variety of methods that explore relationships between genetic variation and environmental gradients (Campbell et al., 2025). GEA analyses are instrumental in understanding the genetic basis of adaptation and can directly inform breeding strategies (Halpin-McCormick et al., 2025). Studies of GEA in crop species have identified major loci for adaptation associated with traits such as drought tolerance and disease resistance (Lasky et al., 2015; Gibson and Moyle, 2020; Lasky et al., 2023), including cold tolerance in wild cranberry subsequently incorporated into breeding efforts (Neyhart et al., 2022) and adaptation to soil conditions in barley (Martin et al., 2025).

Using multiple analytical methods provides an opportunity to best operationalize climate resilience in germplasm collections. These methods can include backcross breeding, population improvement through recurrent selection, or climate matching for targeted germplasm transfer. Different GEA approaches provide information that can be used for plant breeding in complementary ways (reviewed in Campbell et al., 2025). For example, environmental genome wide association studies (EGWAS - Yoder et al., 2014) and F_ST_ outlier and weighted-Z analysis (WZA; Booker et al., 2024) methods can identify specific genes or genomic regions underpinning traits of interest; Focused Identification of Germplasm Strategy (FIGS - Street et al., 2016) and Redundancy Analysis (RDA - De Kort et al., 2014; Rellstab et al., 2021) methods can identify promising populations and analogous environments where they may perform well; and multiple methods (e.g., Todesco et al., 2020) can predict the value of accessions as parents for population improvement (e.g. Halpin-McCormick et al., 2025).

Here we investigated the potential for environmental adaptation in a collection of *Dioscorea* yam accessions. Specifically, we asked (1) how genetic and environmental variation is partitioned among species, hybrid groups and geography in this panel, and whether isolation by environment can be detected after accounting for isolation by distance; (2) whether there are opportunities for allele mining in genomic regions associated with climatic and edaphic variation; and (3) how these patterns can inform breeding, by identifying promising parents and populations at risk under future climates. In this study we applied EGWAS, F_ST_ outlier and WZA scans, RDA, genomic offset and environmental genomic prediction. Together these strategies provide targeted options for university, national and international breeding programs to develop more climate resilient cultivars that can in turn improve food security.

## Materials and Methods

### Data Acquisition

Whole-genome resequencing data for a georeferenced collection of cultivated *D. rotundata* (2n = 2x = 40), wild *D. abyssinica* and semi-wild *D. praehensilis* (both 2n = 2x = 40) and their hybrids were obtained from Scarcelli et al. (2019a, 2019b; https://doi.org/10.5281/zenodo2540773). Reads had been aligned to the *D. rotundata* reference genome TDr96 (Tamiru et al., 2017). The dataset comprised 167 accessions collected at 148 unique sites in Benin, Ghana, Nigeria and Cameroon (**Table 1**) and a VCF file of 3,570,940 SNPs. For each collection site we extracted the 19 WorldClim bioclimatic variables (BIO1–BIO19; Fick & Hijmans, 2017) and 17 soil variables from SoilGrids (ISRIC; Hengl et al., 2017) (**Table S1**). Latitude and longitude were used only as geographic covariates, not as environmental variables. Genotype calls were converted to alternate-allele dosages (0, 1, 2), and genotypes, metadata and environmental values were matched by accession identifier.

#### SNP filtering and missing-data sensitivity

All analyses except the sensitivity analysis described below (population structure, diversity, F_ST_, isolation by distance and environment, EGWAS, WZA, LFMM, BSLMM, RDA, genomic offset and genomic prediction) used biallelic SNPs with no missing calls in any accession (63,142 SNPs; **Figure S1**), so that every analysis used the same genotype matrix without imputation; imputing missing calls in two wild species aligned to the *D. rotundata* reference, without a reference panel, would itself introduce errors. Because the wild species were aligned to the *D. rotundata* reference, missing calls are expected to be more frequent, and non-random, in regions divergent from TDr96, so this filter may under-represent species-specific variation. We therefore calculated, for each genetic group, the proportion of the 3,570,940 SNPs with at least one missing call (**Table S2**) and built a second dataset of biallelic SNPs with ≤ 20% missing calls and minor allele frequency (MAF) ≥0.05 using bcftools (Danecek et al., 2021), imputed with Beagle 5.4 (Browning et al., 2018). Population structure, partial RDA and EGWAS were repeated on this dataset and compared with the main results using Procrustes analysis of PC1–4, the environment-only RDA fraction and the overlap of associated 10-kb windows (**Table S3**).

### Population structure

Population structure was described by principal component analysis (PCA) of SNPs with MAF ≥ 0.05 thinned to one SNP per 7 kb (∼7,200 SNPs), and by spatially explicit ancestry estimation with tess3r (Caye et al., 2016) for K = 2–7, with ancestry coefficients interpolated across West Africa (**Figures 1, S2, S3**). Accessions were assigned to ancestry clusters at K = 7, and the 14 admixed accessions (three hybrid combinations) were pooled into one hybrid group for analysis, giving five analysis groups: *D. rotundata* (n = 80), *D. abyssinica* (n = 29), western *D. praehensilis* (n = 26), Cameroonian *D. praehensilis* (n = 18) and hybrids (n = 14). The 153 non-hybrid accessions were treated as non-admixed, and only these were used in within-species analyses (*D. rotundata*, n = 80; *D. praehensilis*, n = 44). A realized genomic relationship (kinship) matrix was computed with the A.mat function of rrBLUP (Endelman, 2011; VanRaden, 2008), and the first six genetic principal components were used as ancestry covariates where stated.

**Figure 1.**
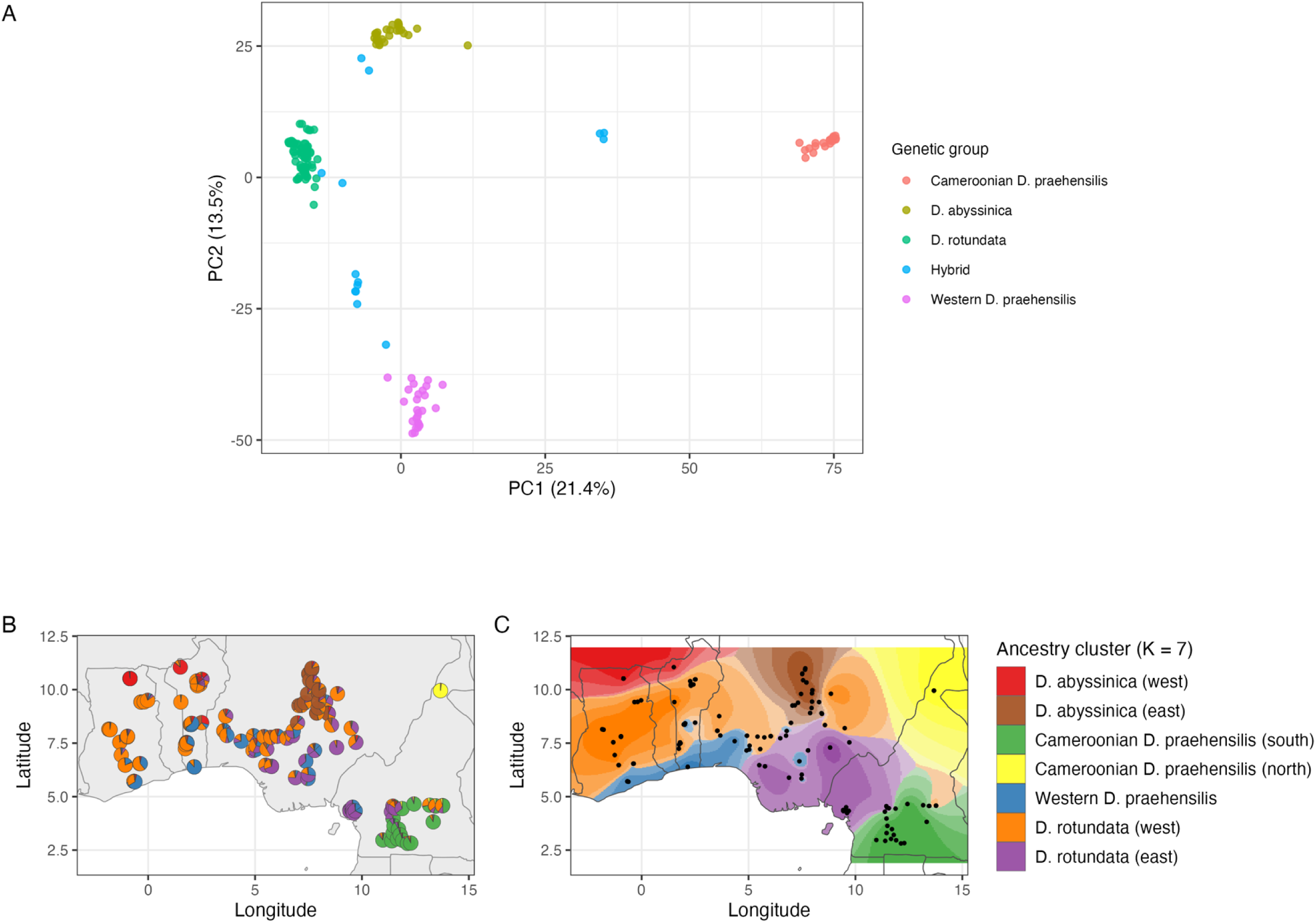
Yam population structure. (A) Principal component analysis of SNPs with MAF ≥ 0.05 thinned to one SNP per 7 kb; accessions are coloured by analysis group, with the three hybrid groups pooled; PC1 and PC2 explain 21.4% and 13.5% of the variance. (B) TESS3 ancestry proportions (K = 7) of each accession, shown as pie charts at its collection site; colours are the seven ancestry clusters, each named after the genetic group most represented among its ten highest-membership accessions (legend); where two clusters share a dominant group, they are distinguished by the location of their highest-membership accessions (west/east or south/north). (C) Spatial interpolation of the TESS3 ancestry coefficients (K = 7) across the sampled range, in the same colours; black points are collection sites. Ancestry for K = 2–7 is shown in Figures S2 and S3.

#### Genetic diversity, runs of homozygosity and private alleles

Within each genetic group we calculated expected heterozygosity at SNP sites, with a sample-size correction, and the folded MAF spectrum (**Table 1; Figure S1**). Runs of homozygosity (ROH) were identified with the consecutive method of detectRUNS (Biscarini et al., 2019), requiring ≥ 20 SNPs and ≥ 1 Mb, gaps between SNPs ≤ 250 kb, and allowing ≤ 1 heterozygous and ≤ 1 missing genotype per run, and F_ROH_ was calculated as the total length of ROH divided by the length of the genome spanned by SNPs (**Figure 2D**). Private alleles were defined as alleles present in one genetic group and absent from all others. Because the number of private alleles increases with sample size and the groups differ greatly in size, we report both raw counts and private allelic richness rarefied to 10 gene copies (Kalinowski, 2004) (**Figure 2B–C**).

**Figure 2.**
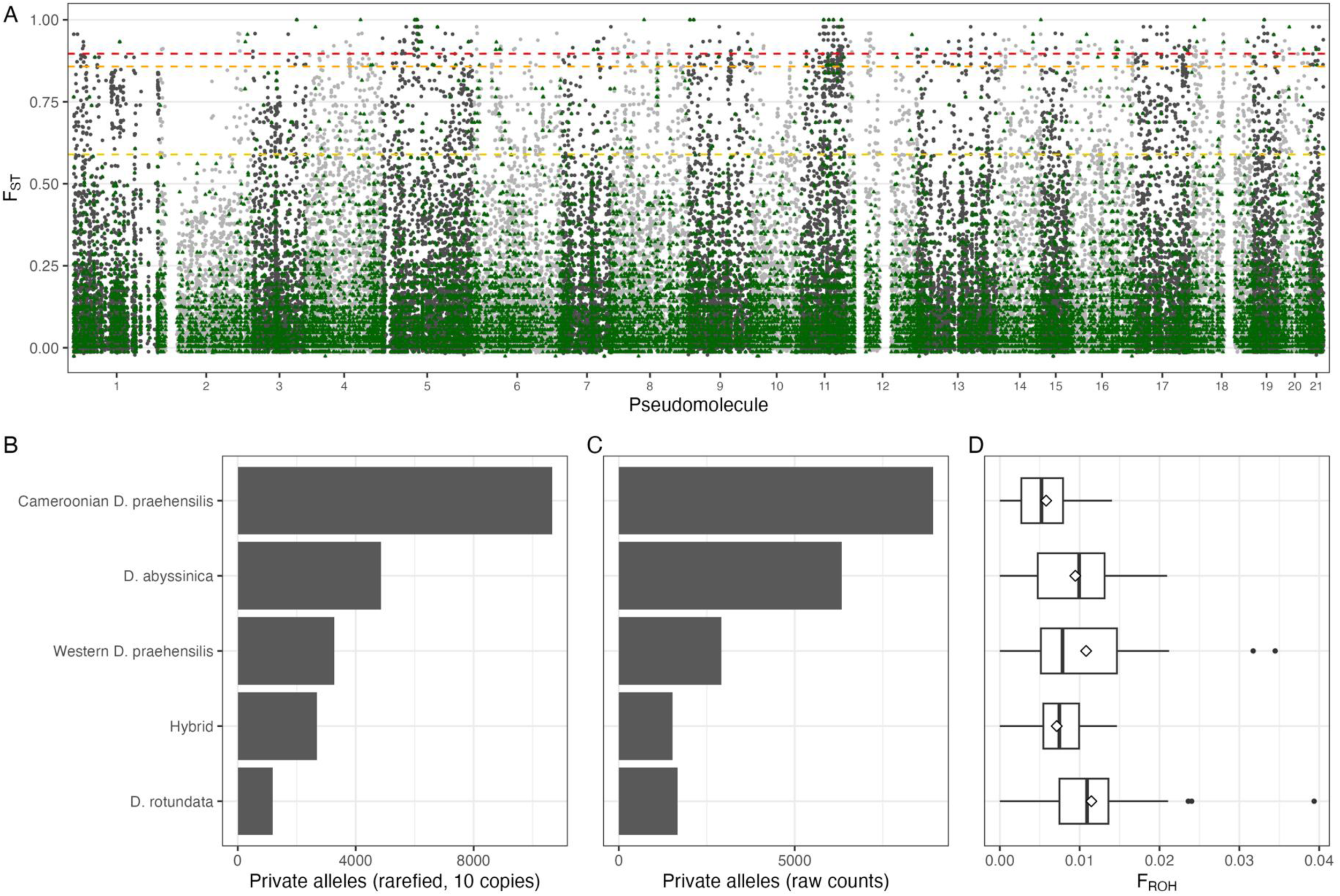
Genetic differentiation, private alleles and inbreeding by genetic group. (A) Per-SNP Weir and Cockerham F_ST_ across the five analysis groups (global contrast) for the 63,142 SNPs, plotted by position along the 21 pseudomolecules (x-axis labels; alternating grey shades); green triangles are SNPs carrying an allele private to one group, and dashed lines are the 95th (yellow, F_ST_ = 0.59), 99th (orange, 0.86) and 99.5th (red, 0.90) percentiles. (B) Private allelic richness rarefied to 10 gene copies and (C) raw counts of private alleles by genetic group. (D) Inbreeding coefficient based on runs of homozygosity of at least 1 Mb (F_ROH_) per accession by genetic group; boxes show the median and interquartile range, whiskers extend to 1.5 × the interquartile range, and white diamonds are group means. Groups are in the same order in B–D. High-F_ST_ loci differentiate groups and may reflect drift, demographic history or population structure as well as selection.

#### Isolation by distance and isolation by environment

To test whether genetic differentiation is associated with environment beyond geographic distance, pairwise genetic distances (Euclidean distances on centered dosages of thinned SNPs) were related to great-circle geographic distance and to environmental distance (Euclidean distance on scaled, VIF-pruned variables; see below) with multiple matrix regression with randomization (MMRR; Wang, 2013) and with partial Mantel tests in vegan (999 permutations), for the full panel and within *D. rotundata*, *D. praehensilis* and *D. abyssinica* (Table S4). The environmental space occupied by each group was summarized by a PCA of the scaled environmental variables (**Figure S4**).

### Genome-environment Association

#### Environmental predictors

All environmental variables were scaled to mean 0 and unit variance. To limit collinearity, variables were removed one at a time, starting with the highest variance inflation factor (VIF), until all VIF < 10; the 16 retained variables (AWCh3, BIO3, BIO8, BIO14, BIO15, BIO18, BIO19, BLDFIE, CECSOL, CRFVOL, OCDENS, OCSTHA, PHIHOX, PHIKCL, SNDPPT and TEXMHT; **Table S1**) were used in RDA, LFMM and isolation-by-environment analyses, whereas EGWAS and genomic prediction analysed all 36 variables individually. The 19 bioclimatic variables were also summarized as three independent components (ICs) with the icaimax function of the R package ica (Bell & Sejnowski, 1995; **Table S5**).

##### F_ST_ outliers and window-based association (WZA)

Per-locus F_ST_ was estimated with the Weir and Cockerham (1984) estimator, computed directly from genotype counts, (i) globally across the five analysis groups (**Figure 2A**) and (ii) for each pair of non-admixed groups. Outliers were defined by the 95th, 99th and 99.5th percentiles of the global per-locus distribution and summarized by chromosome (**Figure S5**), and outliers at private alleles were flagged. Because single-SNP F_ST_ outliers largely reflect divergence among groups, environmental associations were tested with the approaches below. Window-based association was tested with the Weighted-Z Analysis (WZA; Booker et al., 2024): per-SNP p-values from the within-species EGWAS were converted to one-sided Z scores and combined within 10-kb windows with weights p(1 − p), where p is the allele frequency, and window scores were standardized within bins of SNP number per window. Windows in the top 0.1% were retained (**Table S6**).

##### Environmental genome-wide association (EGWAS)

EGWAS was performed with each of the 36 environmental variables and the three ICs as response variables, each rank-based inverse-normal transformed to remove the leverage of extreme values, using the linear mixed model y = Xb + gβ + u + e, with u ∼ N(0, σ²_u_K), where K is the kinship matrix. The model was fitted with the EMMAX/P3D approach (Kang et al., 2010), equivalent to rrBLUP::GWAS with P3D = TRUE: variance components were estimated once per variable by REML (mixed.solve in rrBLUP; residual-to-genetic variance ratio bounded at ≥ 0.05 to avoid boundary fits), and each SNP was then tested by generalized least squares. For the full panel the analysis contained the first six genetic principal components (Q + K model). The analysis was repeated within *D. rotundata* (kinship and two within-species principal components) and within *D. praehensilis* (kinship and a western/Cameroonian lineage covariate), using non-admixed accessions. *D. abyssinica* (n = 29) was not analyzed separately for associations, because power was already negligible within *D. praehensilis* (n = 44; **Figure S7**). In each set, SNPs with MAF ≥ 0.05 were tested (27,785 in the full panel, 20,432 within *D. rotundata* and 31,626 within *D. praehensilis*). For each SNP we report the allelic effect (β per alternate allele, in standard deviations and original units), its standard error, the allele frequency and the proportion of residual variance explained by the SNP, PVE = t²/(t² + df), where t is the SNP test statistic. Significance was assessed at a Benjamini–Hochberg false discovery rate (FDR) of 0.05 (Benjamini & Hochberg, 1995), and the genomic inflation factor (λ) is reported for every variable and set (**Table S7**). Associations with IC1–IC3 are shown in **Figure S6**.

##### Latent factor mixed models

Genotype–environment associations were also tested with latent factor mixed models (lfmm2 in LEA; Frichot et al., 2013; Gain & François, 2021) using the VIF-pruned variables, with K = 7 latent factors for the full panel, matching the K = 7 ancestry clusters, and K = 3 within *D. rotundata* and K = 2 within *D. praehensilis*, chosen from snmf cross-entropy. P-values were calibrated by genomic control, and SNPs significant at FDR 0.05 were compared with the EGWAS results (**Table S8**).

##### Genetic architecture and statistical power

To estimate genetic architecture, we fitted Bayesian sparse linear mixed models (BSLMM; Zhou et al., 2013) in GEMMA v0.98.6 (Zhou & Stephens, 2012) to IC1–IC3, BIO12, BIO17 and PHIHOX, for the full panel and within *D. rotundata* (sample sizes within *D. praehensilis* and *D. abyssinica*, 44 and 29, are too small for reliable estimates of PVE and PGE), using three chains of 10 million sampling steps after 1 million burn-in steps. We report posterior medians and 95% credible intervals for the proportion of variance explained by all SNPs (PVE), the proportion of that variance attributable to sparse, larger-effect SNPs (PGE) and the number of such SNPs (n_γ_) (**Tables S9–S10**). Power to detect a single locus was estimated by simulation on the observed genotypes and kinship: for the full panel and each within-species set we simulated 100 response variables per scenario in which one randomly chosen SNP explained 1, 2, 5, 10 or 20% of the variance against a polygenic background explaining 30% and recorded the proportion detected at FDR 0.05 with the EGWAS model (**Figure S7**).

##### Linkage disequilibrium and candidate genes

Linkage disequilibrium (LD) decay was estimated from r² between 3,000 randomly chosen anchor SNPs and all SNPs within 100 kb on the same chromosome, for the full panel and within *D. rotundata*, using the relaxed, imputed SNP set for marker density. Mean r² in log-spaced distance bins was fitted with r² = c + a·exp(−bd) by nonlinear least squares, and we report the half-decay distance (ln 2/b) and, where reached, the distance at which r² falls to 0.2 (**Figure S8**). Each associated marker received a priority score of 2 if it was detected within a species, plus 1 each if it fell in a top WZA window, was associated with more than one variable, or was also detected by LFMM (possible range 0–5; observed maximum 3; **Tables S11–S12**). Genes were sought near the 223 markers with a score of at least 2; 146 of these had at least one annotated gene within ±10 kb (**Table S12**). Genes near these markers were identified in the TDr96_F1 v1.0 gene annotation of *D. rotundata* (Tamiru et al., 2017), obtained from the Yam Genome Hub (https://yam-genome-hub.southgreen.fr/). All genes overlapping a window of ±10 kb around each marker, about three times the LD half-decay distance (≈ 3.3 kb; **Figure S8**), were retained. The full-panel markers associated with more than one variable that also fell in a top WZA window were annotated in detail: each was classified as lying in a coding exon, intron, untranslated region or intergenic sequence; for coding variants, the codon position and the reference amino acid were recorded. *Arabidopsis thaliana* orthologs were identified as reciprocal best hits (Moreno-Hagelsieb & Latimer, 2008) using BLASTP 2.12.0+ (Camacho et al., 2009; E ≤ 10^−5^) between the TDr96_F1 v1.0 predicted proteins (26,198 sequences; release 20170801) and the Araport11 representative gene models (27,650 proteins; release 20250411; Cheng et al., 2017). A yam and an *Arabidopsis* protein were considered orthologs only when each was the other’s best hit; otherwise the best *Arabidopsis* match is reported as a homolog. Functional descriptions and gene types were taken from Araport11, and protein domains from the InterPro annotation of the TDr96 gene models (Paysan-Lafosse et al., 2023). Because the Araport11 representative protein set excludes transposable-element genes, proteins carrying only transposase domains were not expected to match it.

#### Climate matching

Redundancy analysis (RDA) performed in vegan (Oksanen et al., 2026) was used to partition genetic variation among environment, geography and ancestry. The response matrix was the dosage matrix of SNPs with MAF ≥ 0.05 thinned to one SNP per 7 kb (7,179 SNPs for the full panel). The explanatory sets were the VIF-pruned environmental variables, geography (scaled latitude and longitude) and ancestry (the first six genetic principal components). We fitted a full model with all three sets, a partial RDA (pRDA) of environment conditioned on geography and ancestry, and a variance partitioning (varpart) to estimate pure and shared fractions as adjusted R² (Capblancq & Forester, 2021). The pRDA was tested with 999 permutations for the whole model, for each constrained axis and for each term (marginal effects). pRDA was repeated within *D. rotundata* (ancestry = within-species principal components) within *D. praehensilis* (ancestry = lineage) and within *D. abyssinica* (ancestry = the western/eastern genetic sub-cluster, defined by k-means clustering, k = 2, of the first two within-species principal components; **Figure S10**) (**Table 2**).

Genomic offset was estimated with the genetic.gap function of LEA (Gain & François, 2021; Gain et al., 2023), which uses an LFMM fitted to the observed genotypes (K = 7) to predict the change in adaptive genetic composition required to track climate change at a given location. Predictors were seven bioclimatic variables retained by VIF pruning (VIF < 10) of the 19 bioclimatic variables alone (BIO3, BIO8, BIO9, BIO14, BIO15, BIO18 and BIO19; **Table S1**); soil variables were excluded because no future projections exist. Climate data were taken from WorldClim 2.1 at 10 arc-minutes for current conditions (1970–2000) and from six CMIP6 models (ACCESS-ESM1-5, BCC-CSM2-MR, CanESM5, CNRM-CM6-1, IPSL-CM6A-LR and MIROC6) for SSP5-8.5, 2081–2100. Offsets were computed for each accession, comparing current and future climate at its collection site, and for every land grid cell within 3° of the sampled range, comparing current and future climate in the same cell. Cells whose current climate fell outside the environmental envelope of the sampled sites (ExDet NT1 < 0 or NT2 > 1; Mesgaran et al., 2014) were masked, and the proportion of cells whose future climate fell outside that envelope is reported (ocean cells were excluded). Offsets were averaged across the six models, and the among-model standard deviation is reported as a measure of agreement. Offsets were also computed within *D. rotundata* and *D. praehensilis* (**Table S13**). Köppen–Geiger classifications at 1-km resolution for 1991–2020 and for 2071–2099 under SSP5-8.5 (Beck et al., 2023) were extracted at each collection site. We recorded which sites changed class between the two periods (**Figure 3C–D; Table S13**).

**Figure 3.**
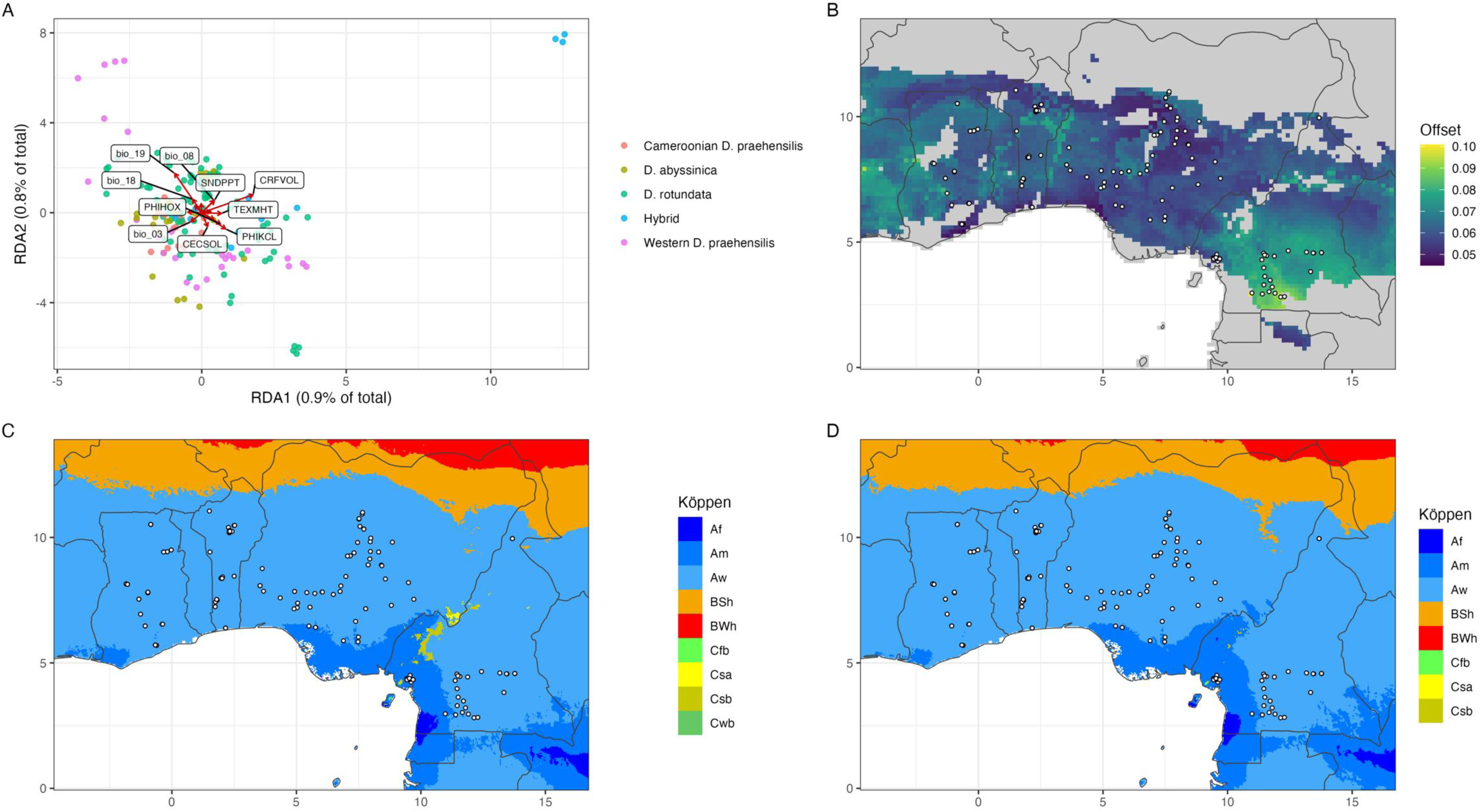
Landscape genomics and climate-change vulnerability. (A) Partial RDA of genetic variation constrained by environment after conditioning on ancestry (six genetic principal components) and geography; points are accessions coloured by genetic group and red arrows are environmental variables; axis percentages (RDA1, 0.9%; RDA2, 0.8%) are percentages of total genetic variance. (B) Mean genomic offset across six CMIP6 models for SSP5-8.5, 2081–2100 (LEA genetic.gap, K = 7); the colour scale runs from low (dark) to high (light); grey cells have current climate outside the environmental envelope of the sampled sites (ExDet), and future climate lies outside that envelope in more than 99.9% of land cells; the ocean is not shown; white points are sampling sites. (C) Köppen–Geiger classes for 1991–2020 and (D) for 2071–2099 under SSP5-8.5 (Beck et al., 2023), with sampling sites in white.

### Parental Selection (Environmental Genomic Selection - EGS)

Genomic prediction was used to estimate genomic estimated adaptive values (GEAVs) for each accession, treating each standardized environmental variable and IC at the collection site as the response, and using all 167 accessions. We compared a structure-only baseline (linear regression on the first six genetic principal components), rrBLUP (ridge regression of SNP effects) and GBLUP with both Gaussian and exponential kernels (K = exp(−(D/θ)²) and K = exp(−D/θ), where D is the scaled Euclidean genetic distance and θ its median), all fitted with rrBLUP (Endelman, 2011). For IC1–IC3, BIO12, BIO17 and PHIHOX we also fitted BayesB and reproducing kernel Hilbert space (RKHS) regression in BGLR (Pérez & de los Campos, 2014; 6,000 iterations, 1,000 burn-in) and random forest (ranger; Wright & Ziegler, 2017). For all 39 variables we also ran GenoBridge v1.0.1 (Singh & Mendu, 2026), a sample-size-adaptive pipeline that selects among ridge regression, random forest, XGBoost, neural-network and stacking models for each variable; with its default settings for this sample size it used 89 genotype principal components, 10-fold cross-validation for model selection and a 20% hold-out set (34 accessions) for accuracy. Because GenoBridge uses its own validation split, its accuracies were compared with our random cross-validation only. Accuracy was determined using the Pearson correlation between out-of-fold predictions and observed values under three cross-validation schemes: random 10-fold (five repetitions); leave-one-group-out, in which each of the five analysis groups is predicted from the others; and spatially blocked, with six blocks defined by k-means clustering of collection coordinates. For each scheme we report the gain in accuracy over the baseline (**Figure S9**). We also evaluated the core-collection training design of the environmental genomic selection framework, in which a core is characterized, and the remaining accessions are predicted. Two 50-accession cores were compared as training sets: the original core and a core re-selected from genotypes only with Core Hunter (Thachuk et al., 2009; **Table S14**); results are reported for the core with the higher mean prediction accuracy (**Table S15**). Models were trained on the core and accuracy was calculated on the 117 non-core accessions. To test whether the core is a better training set than chance, it was compared with 50 random training sets of the same size and 50 random sets stratified to the core’s genetic-group composition, each predicting its own non-training accessions; for every training set we report the pooled correlation, the mean within-group correlation and the gain over the structure-only baseline fitted to the same training set (**Figure S9D–F**). GEAVs used to rank candidate parents were the out-of-fold predictions of the rrBLUP or GBLUP model with the highest spatially blocked accuracy, expressed as deviations from the mean of each accession’s genetic group (**Tables S16–S17; Figure S10**). To display residual structure within groups in **Figure S10**, accessions in each group were ordered by the first principal component of their within-group genotypes (SNPs with within-group MAF ≥ 0.05, thinned to one per 7 kb), and two sub-clusters per group were defined by k-means clustering (k = 2) of the first two within-group principal components.

### Software and code availability

Analyses were run in R v4.4.2 (R Core Team, 2024) using vcfR 1.16.0 (Knaus & Grünwald, 2017), SNPRelate 1.40.0 (Zheng et al., 2012), rrBLUP 4.6.3, vegan 2.7-3, LEA 3.18.0, BGLR 1.1.4, ranger 0.18.0, ica 1.0-3, detectRUNS 0.9.6, corehunter 3.2.3, tess3r, geosphere 1.6-8, terra 1.9-1 and geodata 0.6-9, with figures drawn in ggplot2 4.0.3, together with bcftools, Beagle 5.4, GEMMA 0.98.6, GenoBridge 1.0.1 and BLAST+ 2.12.0. Scripts that reproduce every analysis, table and figure from the input data, with one configuration file recording all settings, are available at https://github.com/kant0063/Yam_GEA.

## Results

### Panel and environmental characterization

The 167 accessions comprised 86 *D. rotundata*, 47 *D. praehensilis* and 34 *D. abyssinica* samples by field identification, collected at 148 unique sites in Benin (n = 36), Ghana (n = 35), Nigeria (n = 55) and Cameroon (n = 41) (**Table 1**). Species and geography were partly confounded as 20 of the 47 *D. praehensilis* were Cameroonian, whereas all but three *D. abyssinica* came from the western part of the species range (Benin, Ghana and Nigeria). Genetic assignment identified 14 hybrid accessions (field-identified as four *D. abyssinica*, six *D. praehensilis* and four *D. rotundata*), and three accessions field-identified as *D. abyssinica* or *D. rotundata* clustered with western *D. praehensilis*. SNPs were distributed across all 21 pseudomolecules (20 chromosomes and one extra scaffold) of the *D. rotundata* reference genome assembly (TDr96; Tamiru et al., 2017) (824–4,522 per pseudomolecule; **Figure S1**). The zero-missing-data filter retained 1.8% of SNPs; per group, 61–86% of the original SNPs had at least one missing call, increasing with group size (86% in *D. rotundata*, n = 80), whereas the per-genotype missing rate was highest in Cameroonian *D. praehensilis* (13.5%) and lowest in *D. rotundata* (3.8%), consistent with reference bias, because reads from the wild species map less fully to the *D. rotundata* genome (**Table S2**). Expected heterozygosity at SNP sites ranged from 0.118 in *D. rotundata*, 0.124 in western *D. praehensilis* and 0.127 in *D. abyssinica* to 0.149 in Cameroonian *D. praehensilis* and 0.153 in hybrids (**Table 1**).

**Table 1.** Composition of the sampling panel, with genetic diversity and environmental range by species. He and environmental ranges refer to the non-admixed genetic group of each species (W, western; C, Cameroonian *D. praehensilis*); totals include hybrids.

| Species (field ID) | Benin | Ghana | Nigeria | Cameroon | Total | Unique sites | Hybrids (K = 7) | He | BIO12 (mm) | Soil pH |
|---|---|---|---|---|---|---|---|---|---|---|
| *D. rotundata* | 19 | 19 | 30 | 18 | 86 | 71 | 4 | 0.118 | 1014–2998 | 5.0–6.5 |
| *D. praehensilis* | 9 | 8 | 10 | 20 | 47 | 46 | 6 | 0.124 (W); 0.149 (C) | 1008–2928 | 5.0–6.5 |
| *D. abyssinica* | 8 | 8 | 15 | 3 | 34 | 33 | 4 | 0.127 | 946–1484 | 5.6–6.4 |
| Total | 36 | 35 | 55 | 41 | 167 | 148 | 14 | 0.118–0.153 | 946–2998 | 5.0–6.5 |

The sampled sites varied in temperature, precipitation and soil variables, but generally reflected conditions suitable for yam production, including mild temperatures (BIO1 range = 21.8–28.2 °C), low-to-medium soil pH (∼5.0–6.5) and relatively high annual precipitation (∼946–2998 mm yr^−1^). Correlations among bioclimatic and soil variables ranged from −0.964 to 0.975 (mean 0.02; **Figures S11–S12**). After VIF pruning, 16 variables were retained (**Table S1**). In an environmental PCA, genetic groups occupied partly distinct environmental space (PC1 46.3%, PC2 16.7%; **Figure S4A**). Cameroonian *D. praehensilis* occupied one end of PC1 (mean score 7.6), at wetter (BIO12 1503–1938 mm), more acidic (pH 5.0–5.6) sites; *D. abyssinica* occupied the other end of PC1 (mean −3.1) at drier (946–1484 mm), less acidic (pH 5.6–6.4) sites; *D. rotundata* and western *D. praehensilis* spanned most of the gradient (**Table 1; Figure S4B**). In MMRR across the full panel, genetic distance increased with both geographic distance (β = 0.15, P = 0.001) and environmental distance (β = 0.34, P = 0.001; R² = 0.20). Within *D. praehensilis*, the environmental effect remained significant (β = 0.14, P = 0.007; geography β = 0.55, P = 0.001; R² = 0.43), but not within *D. rotundata* (β = −0.07, P = 0.25; geography β = 0.23, P = 0.001; R² = 0.04) (**Table S4**). Within *D. abyssinica*, genetic distance was strongly structured by geography (β = 0.86, P = 0.001; R² = 0.75), with no environmental effect (β = 0.02, P = 0.59; partial Mantel r = 0.04, P = 0.26). Isolation by environment was detectable across the panel and within *D. praehensilis*, where it is partly confounded with the western–Cameroonian split. The genetic structure within cultivated *D. rotundata* and *D. abyssinica* follows geography.

### Population structure

A principal component analysis was conducted to explore genetic variation among five analysis groups: four non-admixed groups (*D. abyssinica*, western *D. praehensilis*, Cameroonian *D. praehensilis* and *D. rotundata*) and the 14 hybrids, which represent three hybrid combinations (*D. abyssinica* × *D. rotundata*, western *D. praehensilis* × *D. rotundata* and Cameroonian *D. praehensilis* × *D. abyssinica*). The first two principal components (PC1 and PC2) explained 21.4% and 13.5% of the total variance, respectively. Distinct clustering among the species was observed (**Figure 1A**). Cameroonian *D. praehensilis*, *D. abyssinica*, *D. rotundata* and western *D. praehensilis* each formed a discrete, compact cluster, with hybrids (all combinations represented by one color) being more dispersed, partially overlapping with multiple species but typically about halfway between their two parent species. To further understand the geographic distribution we explored spatial interpolation of the population structure (K = 2–7) across the sampled range (**Figures S2–S3**). At lower K values (K = 2–3), individuals clustered into broad genetic groups corresponding to the t sampled *Dioscorea* species, but as K increased (K = 4–6), additional substructure and admixture patterns became evident among populations in areas of geographic overlap. At K = 7, the clusters followed both genetic group and geography, with *D. rotundata* and *D. abyssinica* spliting into a western and an eastern cluster, Cameroonian *D. praehensilis* into a southern and a northern cluster (the latter concentrated at the north-eastern edge of the range), and western *D. praehensilis* formed a single cluster; accessions with mixed ancestry occurred where these clusters met (**Figure 1B–C**).

### Genome Environment Association - Potential for allele mining

#### F_ST_ Outlier Analysis and Latent Factor Mixed-Model

Private alleles were present in all genetic groups. Cameroonian *D. praehensilis* had the most private alleles (8,934 counted directly; 10,658 after rarefaction to 10 gene copies) and *D. rotundata* the fewest (1,668 counted directly; 1,179 rarefied). Rarefaction removes the effect of sample size: it lowers the count for large groups such as *D. rotundata* (n = 80) and can raise it for small groups, because alleles that are rare in other groups become private once every group is reduced to the same number of gene copies. Hybrids ranked lowest on direct counts (1,529) but above *D. rotundata* after rarefaction (2,681), so their apparent scarcity of private alleles reflected sample size (Figure 2B–C). When exploring the overall levels of inbreeding across the species we found that F_ROH_ based on runs of homozygosity of at least 1 Mb was very low (group means 0.006 in Cameroonian *D. praehensilis* to 0.011 in *D. rotundata*; maximum 0.039), indicating little inbreeding in the *Dioscorea* yams that make up the agriculturally relevant species complex (**Figure 2D**). Per-SNP Weir and Cockerham F_ST_ across the five analysis groups was highest at loci differentiating groups (pairwise genome-wide F_ST_ 0.15–0.37, highest between Cameroonian *D. praehensilis* and the remaining groups), and 346 of 3,155 outliers (11%) were at private alleles (**Figure 2A**). There was not an even distribution of outlier markers across the genome, for example, the chromosome with the most outlier SNPs was pseudomolecule 11 (n = 318) and the one with the least was pseudomolecule 21 (n = 47) (**Figure S5**). Outliers tracked group membership and hybrid status rather than environmental gradients, likely meaning that they are markers of divergence among groups, which may reflect drift, demographic history or selection, rather than local adaptation. Within-species WZA identified 214 distinct windows in the top 0.1% (across variables) in *D. rotundata* and 286 in *D. praehensilis*, with 2 and 53 window–variable combinations overlapping with EGWAS associations (**Table S6**). LFMM identified 333 SNPs associated with at least one environmental variable at FDR 0.05 in the full panel (K = 7), 9 within *D. rotundata* (K = 3) and 1,989 within *D. praehensilis* (K = 2); the large number in *D. praehensilis* most likely reflects the confounding of environment with the western– Cameroonian split rather than local adaptation; only one LFMM SNP (full panel) was also detected by EGWAS (**Table S8**).

#### Environmental Genome Wide Association

Across the full panel, the EGWAS using 19 bioclimatic variables and 17 soil variables identified 19 marker–environment associations at 16 unique markers (**Figure 4**). Including kinship and six genetic principal components as covariates reduced the likelihood of detecting SNPs only associated with background genetic relatedness (Yoder et al. 2014; Anderson et al. 2016; Bandillo et al. 2017; Lei et al. 2019). Significant markers were identified on 9 of the 21 pseudomolecules and were detected for 9 of the 36 precipitation, soil and temperature variables, most often precipitation of the wettest quarter (BIO16; 7 markers) and of the coldest quarter (BIO19; 4 markers). Three of the 16 markers (19%) were associated with multiple variables (**Table S18; Figures S13–S15**), consistent with pleiotropy, although because many variables are correlated (**Figures S11–S12**), associations of one marker with several variables may reflect a single underlying gradient. We summarized the 19 bioclimatic variables using Independent Component analysis (ICA) to reduce multicollinearity and capture independent climatic gradients (**Table S5**). Independent Component analysis (ICA) created independent factors for the climate variables with IC1 being related to temperature (BIO1, BIO8, BIO9, BIO11), IC2 made up of Precipitation (BIO12, BIO13, BIO16, BIO19), and IC3 being a mix of Seasonality (BIO4, BIO15) and Temperature range (BIO2, BIO7) (**Table S5**). Independent Components 1–3 were uncorrelated (pearson correlation r = 0), confirming they capture independent axes of climatic variation. The same mixed model identified no SNPs significantly associated with IC1, IC2 or IC3 (λ = 0.92, 0.93 and 0.97; **Figure S6**). Among markers that were associated with more than one climatic variable it was more common that they were associated with water stress (**Figure S16**). Two of the three multi-variable markers also fell in top WZA windows and were annotated in detail (**Table S12**). There were only weak geographic differences in allelic state for the three multi-variable markers associated with environmental variation (**Figure S17**), with their alternative alleles occurred mainly in the western and central part of the range and were rare or absent in Cameroon, so allele states also tracked genetic group.

**Figure 4.**
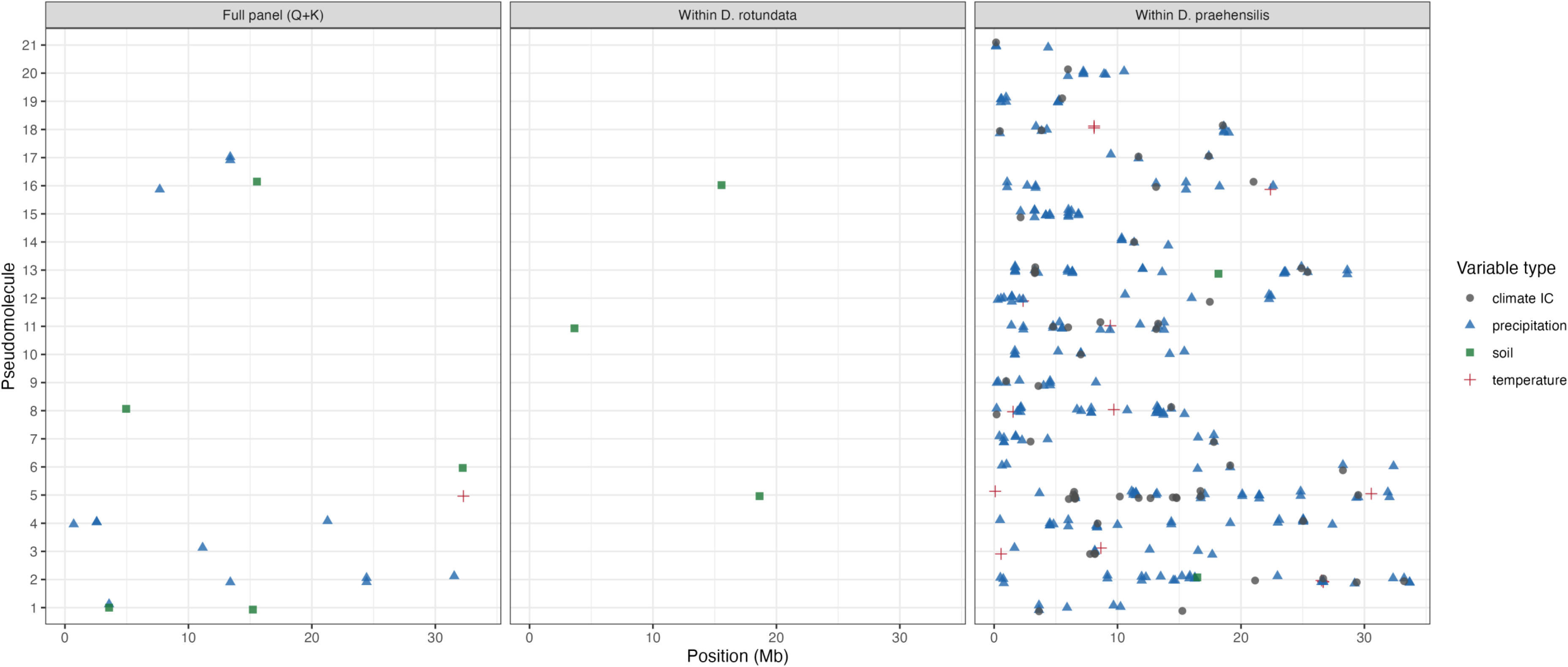
Significant marker–environment associations (FDR 0.05) by pseudomolecule (y-axis) and position (Mb) for the full panel (Q + K model) and within *D. rotundata* and *D. praehensilis*. Colours and shapes show the type of variable (precipitation, temperature, soil, climate IC). Overlapping symbols mark markers associated with more than one variable. The full panel yielded 19 associations at 16 unique markers, *D. rotundata* three and *D. praehensilis* 327 (the latter largely reflecting the western–Cameroonian split). BSLMM estimates of genetic architecture are given in Table S9.

Two of the three multi-variable markers, BDMI01000002.1_24434489 and BDMI01000017.1_13385808, also fell in top WZA windows and were annotated in detail. Each lies within a single annotated gene, and no other gene occurs within 10 kb (the nearest flanking genes are 17.8–46.9 kb away; **Table S12**). Marker BDMI01000002.1_24434489 is in the second coding exon of Dr02609 (BDMI01000002.1:24,433,752–24,436,488). The alternative allele changes the second position of codon 171 and therefore replaces isoleucine with asparagine or lysine; this residue is isoleucine or valine in the homologous yam proteins Dr07933 (91% identity), Dr02608 and Dr09098. Protein Dr02609 has no annotated InterPro domain. Its best *Arabidopsis* match, AT5G03990 (29% identity over 230 aligned residues; E = 5 × 10^−24^), and the second match, AT3G51940 (E = 9 × 10^−17^), are protein-coding genes without a functional description in Araport11, with AT5G03990 matching Dr09098 most closely. Protein Dr02609 belongs to a small yam gene family that includes the neighboring gene Dr02608, 31.6 kb away, so it has no one-to-one *Arabidopsis* ortholog and no predicted function. BDMI01000017.1_13385808 lies in the fourth intron of Dr01303 (BDMI01000017.1:13,377,427–13,389), whose only annotation is a plant Ptta/En/Spm transposase domain (IPR004252), characteristic of CACTA DNA transposons (Wicker et al., 2007). Protein Dr01303 had no significant match among Araport11 protein-coding genes (best E = 3.4), shares 58–75% identity with several other yam genes, and is one of 161 TDr96 gene models carrying this domain; it is therefore most likely part of a transposable element. Neither marker thus implicates a gene of known function. Dr02609, carrying a missense variant, is the more plausible candidate, but both associations are better treated as targets for fine mapping than as evidence for a specific adaptive mechanism.

Within *D. rotundata*, EGWAS identified three associations at three markers, two with soil pH (PHIHOX) and one with cation exchange capacity (CECSOL). One of the markers associated with soil pH, BDMI01000016.1_15551415, was the only significant full-panel marker that was also recovered within species. Within *D. praehensilis*, 327 associations at 218 markers were detected, almost all with precipitation variables and IC3; seasonality and temperature range), but the variance-component estimates reached their upper bound for 34 of the 39 variables and genomic inflation ranged down to λ = 0.62, so these associations most likely reflect the western–Cameroonian split rather than local adaptation and are not interpreted further. Genomic inflation ranged from λ = 0.84 to 1.09 in the full panel and from 0.92 to 1.08 within *D. rotundata* (**Table S18**). Individually significant markers explained 12–16% of the residual variance in the full panel (median 14%) and 24–28% within *D. rotundata* (**Table S18**).

In BSLMM, SNPs jointly explained most of the variation in every variable tested (full panel: IC1 97%, 95% credible interval 84–100%; IC2 70%, 46–97%; IC3 96%, 83–100%; BIO12 95%; BIO17 88%; PHIHOX 97%), reflecting how strongly environment of origin is predicted by relatedness. For annual precipitation (BIO12; PGE = 0.90, 0.58–1.00; n_γ_ = 24), dry-quarter precipitation (BIO17; PGE = 0.82, 0.33–0.99; n_γ_ = 18), soil pH (PHIHOX; PGE = 0.77, 0.12–0.99; n_γ_ = 32) and IC2 (PGE = 0.87; n_γ_ = 12), most of this variance was attributed to a modest number of sparse, larger-effect loci, whereas for IC1 and IC3, PGE was lower and highly uncertain (0.41 and 0.33, credible intervals spanning 0–0.97). The same pattern held within *D. rotundata* (e.g. BIO12 PGE = 0.83, 0.71–0.94; n_γ_ = 7) (**Table S9**). Power simulations showed that even a locus explaining 20% of variance would be detected at FDR 0.05 with probability 0.60 in the full panel, 0.19 within *D. rotundata* and 0.01 within *D. praehensilis* (association scans and power simulations were not run within *D. abyssinica*, n = 29), and a locus explaining 10% with probability 0.16, 0.02 and 0.00 (**Figure S7**); loci of small to moderate effect are therefore unlikely to be detected individually.

#### Climate Matching - Finding places where current populations may work

A redundancy analysis (RDA) was conducted to assess the extent to which environmental variables explain genomic variation among the studied *Dioscorea* populations. There were substantial relationships between the environmental and soil variables (**Figure S12**), in particular soil and precipitation variables (**Figures S11–S12**). In the full model, environment, geography and ancestry together explained 57.4% of genetic variance (adjusted R² = 0.50), and the first two axes respectively explained 21.2% (RDA1) and 13.4% (RDA2) of total genetic variance (full-model ordination not shown). Clustering patterns among populations showed genetic differentiation consistent with species and geographic distribution. Thus, these axes likely reflect species divergence rather than environment. In the full model, *D. rotundata* was associated with soil texture and pH (TEXMHT, SNDPPT, PHIHOX, PHIKCL and AWCh3); Cameroonian *D. praehensilis* with isothermality (BIO3), precipitation of the driest and warmest quarters (BIO14, BIO18) and soil organic carbon density (OCDENS); and *D. abyssinica* with precipitation seasonality (BIO15) and soil organic carbon stock (OCSTHA). In contrast, western *D. praehensilis* separated mainly along an ancestry axis rather than an environmental one.

In the partial RDA (pRDA), which conditions environment on ancestry and geography (the full model, by contrast, includes all three sets as explanatory variables), environment explained 1.8% of genetic variance (adjusted R²; P = 0.001), compared with 25.0% for ancestry alone and 0.4% for geography alone; 23.0% was shared, mostly between environment and ancestry (**Table 2**). In the pRDA ordination (**Figure 3A**), the first two constrained axes explained 0.9% and 0.8% of total genetic variance, accessions of all groups largely overlapped except a few hybrid and *D. rotundata* accessions, and coarse fragment volume (CRFVOL) and precipitation of the coldest quarter (BIO19) had the longest arrows; once species and geography are accounted for, the remaining environment-associated variation therefore relates mainly to soil texture and to dry-season rainfall (in West Africa the coldest quarter falls in the dry season). Within *D. rotundata*, environment conditioned on ancestry and geography explained 1.1% (P = 0.026), and within *D. praehensilis* 2.4% (P = 0.001). Within *D. abyssinica*, environment conditioned on geography and the western/eastern sub-cluster explained 3.6% (P = 0.002), but only one constrained axis and no individual environmental variable was significant; the sub-cluster and geography were almost entirely collinear, and with 29 accessions and 12 environmental variables this estimate should be treated cautiously (full-model R² = 0.65, adjusted R² = 0.25). In the full panel, four of 16 constrained axes and 11 environmental terms were significant by permutation. Three soil physical variables (available water capacity, AWCh3; coarse fragment volume, CRFVOL; texture class, TEXMHT), three soil chemical variables (cation exchange capacity, CECSOL; organic carbon density, OCDENS; pH, PHIHOX), isothermality (BIO3) and four precipitation variables (BIO14, BIO15, BIO18, BIO19). Within *D. rotundata*, isothermality and two soil variables (OCDENS, TEXMHT) remained significant, and within *D. praehensilis* two temperature variables (BIO3, BIO8), two precipitation variables (BIO15, BIO19) and four soil variables (BLDFIE, CECSOL, OCDENS, OCSTHA) (**Table 2**). Only a small fraction of genetic variation can therefore be attributed to environment independently of species and geography; the variables significant in the full panel and within both *D. rotundata* and *D. praehensilis*, isothermality (BIO3) and soil organic carbon density (OCDENS), are the most credible targets for breeding.

**Table 2.** Variance partitioning of genetic variation by redundancy analysis for the full panel and within the non-admixed *D. rotundata*, *D. praehensilis* and *D. abyssinica* accessions. Ancestry is represented by the first six genetic principal components in the full panel, by within-species principal components in *D. rotundata*, by the western/Cameroonian lineage in *D. praehensilis* and by the western/eastern genetic sub-cluster in *D. abyssinica*. Fractions are adjusted R² (%) from variance partitioning among environment (VIF-pruned variables), ancestry and geography (latitude and longitude); ∩ denotes variance shared only by the sets named. P for the environment-only fraction is from 999 permutations of the partial RDA (pRDA) conditioned on ancestry and geography; axes and terms (marginal effects) were tested with 999 permutations at P < 0.05.

|  | **Full panel** | *D. rotundata* | *D. praehensilis* | *D. abyssinica* |
|---|---|---|---|---|
| Accessions | 167 | 80 | 44 | 29 |
| SNPs (MAF ≥ 0.05, thinned) | 7,179 | 4,999 | 8,372 | 5,893 |
| Environmental variables (VIF < 10) | 16 | 13 | 12 | 12 |
| Ancestry covariates | 6 genetic PCs | 6 within-species PCs | Lineage | Western/eastern sub-cluster (1) |
| Full model R² (adjusted R²) | 0.574 (0.502) | 0.482 (0.295) | 0.685 (0.516) | 0.651 (0.249) |
| Environment only (P) | 1.8 (0.001) | 1.1 (0.026) | 2.4 (0.001) | 3.6 (0.002) |
| Ancestry only | 25.0 | 21.4 | 1.2 | –0.2 |
| Geography only | 0.4 | 0.3 | 0.6 | 0.03 |
| Environment ∩ ancestry | 11.0 | 1.6 | 12.0 | 1.6 |
| Ancestry ∩ geography | 0.6 | 0.6 | 3.2 | 4.5 |
| Environment ∩ geography | 0.4 | 0.3 | 3.5 | 2.8 |
| Shared by all three | 11.0 | 4.3 | 28.7 | 12.6 |
| Residual | 49.8 | 70.5 | 48.4 | 75.1 |
| Significant pRDA axes | 4 of 16 | 0 of 13 | 3 of 12 | 1 of 12 |
| Significant environmental terms | AWCh3, BIO3, BIO14, BIO15, BIO18, BIO19, CECSOL, CRFVOL, OCDENS, PHIHOX, TEXMHT | BIO3, OCDENS, TEXMHT | BIO3, BIO8, BIO15, BIO19, BLDFIE, CECSOL, OCDENS, OCSTHA | none (closest: CECSOL, P = 0.050) |

Per-accession genomic offset for SSP5-8.5, 2081–2100 ranged from 0.048 to 0.101 (median 0.062) and was highest for Cameroonian *D. praehensilis* (median 0.079, compared with 0.060–0.062 in the other groups) (**Table S13**). Offset declined with latitude (P < 0.001; R² = 0.19), but this was driven entirely by the southern Cameroonian sites; without these accessions, there was no trend (P = 0.90). Agreement among the six climate models was moderate (median among-model SD = 46% of the mean offset). Across the mapped region, offsets ranged from 0.045 to 0.101 (median 0.062) in cells whose current climate lay within the sampled region (52% of 8,308 land cells). The highest values were found in southern Cameroon and toward the western edge of the range and lowest across the interior of Nigeria and Benin. Future climate fell outside the sampled envelope in more than 99.9% of land cells (**Figure 3B**). Only one of the 148 sites changed Köppen–Geiger class between 1991–2020 and 2071–2099 (tropical savanna, Aw, to tropical monsoon, Am, at a Cameroonian *D. praehensilis* site), so no statistical comparison with offsets was possible (**Figure 3C–D; Table S13**). The shift in climate at each site is shown in **Figure S18**.

#### Parental selection – environmental genomic prediction

As expected, prediction accuracy varied based on the cross-validation scheme (**Figure S9**). Random 10-fold cross-validation had a mean accuracy across variables was 0.69–0.73 for the genomic models, but the structure-only baseline reached 0.61, so genomic models added only 0.08–0.11. Using Leave-one-group-out cross-validation caused accuracy to fall to 0.05–0.23 (baseline −0.08; gain 0.13–0.28), and spatially blocked cross-validation (0.01–0.08) was no better than the baseline (gain −0.06 to 0.04). Most of the predictive ability under random cross-validation therefore reflected population structure, and predictions did not transfer to new geographic areas. Model class had little influence as BayesB and RKHS were marginally best under random and leave-one-group-out cross-validation (**Figure S9**). GenoBridge selected ridge regression for 29 of the 39 variables and the stacking ensemble for the other 10. Its hold-out accuracy averaged 0.70 (median 0.74; range 0.02 for CECSOL to 0.86 for IC3), the same as the best of our models under random cross-validation (mean 0.70, median 0.73) and 0.09 above the structure-only baseline; GenoBridge was more accurate for 22 variables and less accurate for 17, with the largest differences for BIO6 (+0.16), IC2 (+0.15) and CRFVOL (−0.26). For the variables fitted with every model (the three climate components IC1–IC3 and the variables most relevant to yam production: annual precipitation, BIO12; dry-quarter precipitation, BIO17; and soil pH, PHIHOX) it averaged 0.79, compared with 0.74 for our best models (**Table S19**). Because GenoBridge’s hold-out set is small (34 accessions) and, like random cross-validation, not separated by group or space, these accuracies are imprecise and partly reflect population structure. Of the two cores, the core re-selected from genotypes gave much higher accuracy than the original core (mean r on non-core accessions 0.62 vs 0.22; within-group r 0.48 vs 0.06; **Table S15**); the original core contained no Cameroonian *D. praehensilis*, so all 18 accessions of that group had to be predicted by extrapolation. Trained on the genotype-based core, the genomic models reached mean accuracies of 0.61–0.62 (gain over the baseline 0.04–0.05), similar to size-matched random (0.62–0.63) and group-stratified (0.61–0.62) training sets, but with higher within-group accuracy (0.46–0.50 vs 0.37–0.40); the core exceeded the random-set mean for 55% of variable–model combinations (**Figure S9D–F**). A core selected from genotypes is therefore as efficient as a random training set of the same size overall and ranks accessions within groups somewhat better, whereas a core lacking a whole genetic group is not. Ranked on within-group, out-of-fold GEAVs, the top candidates for tolerance to low soil pH were all *D. rotundata*: CR5387, CR5390 and CR5392 from Cameroon and CR5574 and CR5523 from Nigeria. For high annual precipitation (BIO12), they were three *D. rotundata* accessions (CR5602 from Nigeria and CR5387 and CR5392 from Cameroon) and two western *D. praehensilis* accessions from Cameroon (P5483 and CR5467) (**Figure S10; Tables S16–S17**). Because spatially blocked accuracy was near zero, these rankings are hypotheses for field evaluation rather than reliable predictions. Within *D. abyssinica*, the two genetic sub-clusters shown in **Figure S10**, which have opposite within-group GEAVs for the temperature variables (BIO1–BIO11), are geographically separated: one comprises the 13 accessions from Ghana and western Benin (0.9°W–2.5°E) and the other the 15 Nigerian accessions and one from eastern Benin (2.5–8.4°E), at overlapping latitudes (8.4–11.1°N). The first within-group principal component explained 22.5% of genetic variation in *D. abyssinica*. These GEAVs therefore reflect a west–east genetic split that coincides with longitude and should not be interpreted as adaptation to temperature within the species.

## Discussion

Using multiple techniques to explore the genetic basis of climate resilience in yam species commonly grown across West Africa revealed substantial genetic diversity, most of it structured by species and geography. Environment, geography and ancestry together explained 57% of genetic variation (adjusted R² = 0.50), but only 1.8% was attributable to environment independently of species identity and geography (**Table 2**); this smaller fraction is the more defensible estimate of the role of local selective pressures in shaping yam genomic diversity (Fournier-Level et al., 2011, Rellstab et al., 2015). BSLMM analysis showed variance in precipitation and soil pH was likely due to a small number of larger-effect loci, however, credible intervals were wide and the sample size was small. This suggests that population improvement (i.e. recurrent selection that accumulates many favorable alleles of small effect at once), is the most promising breeding strategy for most variables. A handful of markers had individually detectable effects (12–28% of residual variance); such loci could instead be tracked with markers and introgressed into elite cultivars by marker-assisted backcrossing, which is efficient only when a few loci of large effect are targeted, although effects estimated for markers detected at low power are likely to be overestimated. This combined genetic architecture provides multiple different potential breeding strategies. Breeding efforts may include many common practices including ennoblement (farmer selections that can be hybrids or wild collections) and targeted intra- and interspecific crosses (Chair et al., 2010). Integrating these practices directly by picking the best potential parents will help improve crop resiliency.

We identified a similar population structure to that previously identified (Scarcelli et al., 2019a) and results from admixture analyses confirmed the documented history of wide hybridization previously identified in yam germplasm collections and documented in yam breeding programs (Mondo et al., 2022). The GEA analyses were consistent with a mixed genetic architecture for environmental adaptation in yam, with some biophysical and bioclimatic traits associated with moderate-large effect loci. As in other GEA studies of structured germplasm (Lasky et al., 2015; Gibson & Moyle, 2020), F_ST_ outliers mainly marked divergence among genetic groups rather than environmental adaptation. For annual precipitation (BIO12), precipitation of the driest quarter (BIO17) and soil pH (PHIHOX), BSLMM attributed most SNP-explained variance to a modest number of larger-effect loci (PGE 0.77–0.90; **Table S9**), although these loci were rarely detectable individually, similar to what has been observed in other crops (Lasky et al., 2015, Exposito-Alonso et al., 2019). In yam production regions, increasing temperature and soil degradation (specifically low soil organic carbon and low pH) are reducing crop yields making climate adapatation a priority for breeding (Mondo et al., 2024; Kouakou et al., 2025). Here we identified markers associated with soil pH at collection sites, including one detected both in the full panel and within *D. rotundata* (**Table S18**); whether they confer tolerance to low pH remains to be tested. In addition, there were moderate correlations between precipitation and soil characteristics (**Figure S12**). There were also accessions that had promising GEAVs for soil conditions making this a reasonable breeding target. Parental selection for climate adaptation can be informed by GEAVs, which are analogous to genomic estimated breeding values (GEBVs) but quantify adaptation to environmental gradients rather than agronomic traits. Because GEAVs were expressed relative to each accession’s genetic group, the candidate parents we report (e.g. CR5387, CR5392 and CR5602) differ from their group mates in predicted adaptation rather than simply reflecting species or region of origin. However, genomic prediction did not transfer across space (spatially blocked r ≈ 0), so these candidates require field evaluation before use as parents.

None of the associations identified here points to a gene of known function. The marker associated with precipitation of the wettest and coldest quarters (BIO16 and BIO19) causes an isoleucine-to-asparagine or -lysine substitution in Dr02609, a member of a small yam-specific gene family without an *Arabidopsis* ortholog or recognized protein domain; the marker associated with BIO13 and BIO16 lies in an intron of Dr01303, which most likely belongs to a CACTA transposon. The latter is consistent with reports that transposable-element variation contributes to climatic adaptation in wild grapevine (Ma et al., 2026) and suggests that future studies in yam should genotype structural and transposable-element variation. Genes within 10 kb of the two within-*D. rotundata* markers that fell in top WZA windows (soil pH on BDMI01000005.1 and cation exchange capacity on BDMI01000011.1) (**Table S12**). Given the low power and the number of tests, we highlight these genes as targets for fine mapping and functional validation rather than established adaptive loci. Two Cameroonian *D. rotundata* accessions, CR5387 and CR5392, ranked among the top five for both tolerance to low soil pH and high annual precipitation, with within-group GEAVs about 1.1–1.2 standard deviations beyond their group mean for each (**Table S17**), making them the most promising candidate parents for wet, acidic environments; CR5390, CR5574 and CR5523 (low pH) and CR5602 (high precipitation) are strong alternatives for a single target, and two western *D. praehensilis* accessions from Cameroon, P5483 and CR5467, could contribute high-precipitation adaptation through interspecific crosses. Because accuracy under random cross-validation largely reflected population structure, accuracy for an unseen genetic group was modest (leave-one-group-out r = 0.05–0.23) and predictions did not transfer to new geographic areas (spatially blocked r ≈ 0; **Figure S9**), these rankings are most reliable for choosing among accessions within the sampled groups and regions, and should be confirmed in field trials before the accessions are used as parents.

In the full RDA model, species separated along climatic and edaphic axes, but these axes largely reflect species divergence (**Figure 3A**; **Table 2**). Isothermality (BIO3) and soil organic carbon density (OCDENS) were significant in the full panel and within both *D. rotundata* and *D. praehensilis*. Projected genomic offsets varied across the region (**Figure 3B**), with moderate agreement among climate models. Offsets were highest for the Cameroonian lineage of *D. praehensilis*, at the wettest, most acidic sites, whereas Köppen–Geiger class changed at only one sampling site (**Figure 3C–D**). The Cameroonian lineage may therefore face the greatest adaptive mismatch as future climates fall outside the sampled range, however, genomic offsets are difficult to interpret as measures of vulnerability (Lotterhos, 2024). If confirmed, the Cameroonian lineage of *D. praehensilis* would be a priority for conservation and germplasm collection. Since associated markers implicate a gene of known function, functional validation is essential before these associations are interpreted as adaptive (Rellstab et al., 2015). There are also potential agronomic adaptations to try to address changing climate, including altering the cropping calendar (Cornet et al., 2025). Two caveats apply to these projections, first, the clonality of cultivated *D. rotundata* suggests that movement by farmers may mean collection sites may not represent the environments in which its alleles were selected. Second, genomic offset assumes that the allele–environment relationships identified by GEA are adaptive and will hold under future climates (Lotterhos, 2024), which we tried to address by masking predictions outside the sampled environmental envelope.

Although environment explained only a small fraction of genetic variation in this panel, genome-environment association analyses can identify useful variation, in particular for abiotic stresses (Gao et al., 2023). Wild relatives of crops have long been exploited by breeders as a source of favorable genetic variation for different traits, including through interspecific crosses (Hajjar and Hodgkin 2007). The domestication history of yam suggests that it may be possible to use wide hybridization more effectively without a large amount of linkage drag (Scarcelli et al., 2017). Successful utilization of abiotic stress tolerance loci could address limitations in expanding yam cultivation. Three of the 16 full-panel markers were associated with more than one variable; because these variables are correlated, this may reflect a shared gradient rather than pleiotropy, and apparent pleiotropy of this kind is common in GEA studies of other crops (Neyhart et al., 2022; Wang et al., 2023; Halpin-McCormick et al., 2025). Because wild and cultivated yams are widely adapted and readily available in genebanks, these genetic resources could be used more extensively than they are at present. There is substantial variation in cultivated yam, the use of marker-assisted backcrossing may provide many benefits especially since some markers have individually detectable effects (12–28% of residual variance; **Table S18**), notably the soil-pH marker BDMI01000016.1_15551415 detected both in the full panel and within *D. rotundata*. A major next step will be to conduct validation experiments by growing clones of specific accessions under the stressed conditions in both field and greenhouse conditions to examine if these effects are as large as predicted. There may be an option to do early-generation selection as there is much overlap between wild, semi-wild, and elite material. The candidate environments and accessions identified here could also guide the Focused Identification of Germplasm Strategy (FIGS; Street et al., 2016) when choosing subsets of the collection for phenotypic screening. Because yam is propagated clonally, heterozygous genotypes are fixed in cultivars, so the effects of heterozygosity at candidate loci should be examined in these validation trials, and scaling up the multiplication of high-quality seed tubers of selected material will be needed to deliver any gains to farmers (Mondo et al., 2024; Cornet et al., 2025).

Our results can be compared with recent landscape-genomic studies of crops and their wild relatives that illustrate both the potential and the limitations of approaches to climatic adaptation. The genetic basis of climatic adaptation is architecturally diverse, and this diversity shapes how candidate loci should be interpreted and used in breeding. For example, in alfalfa and *Medicago truncatula*, GEA candidates were found in highly conserved genes, and a subset were also associated with phenotypic traits, consistent with a largely polygenic basis for climatic and edaphic adaptation (Blanco-Pastor et al., 2021). Structural and transposable-element variants can also be prominent, as in wild apple (*Malus baccata*), where both SNPs and structural variants were associated with climate, with northeastern genetic clusters projected to face the highest risk of maladaptation (Su et al., 2026), while in wild Amur grape (*Vitis amurensis*) transposable-element variants were specifically associated with climatic adaptation (Ma et al., 2026). In *Populus species,* using multiple GEA approaches (e.g. genomic offset analysis, GWAS, and genomic prediction) identified signals of adaptation and for parental identification for breeding (Fitzpatrick & Keller, 2015; Xiang et al., 2024; Guo et al., 2025). Multiple approaches were also used in North American wild Vitis species and worked best to identify accessions that were adapted to extreme climates and of use to grapevine breeding (Aguirre-Liguori et al., 2022).

The *D. praehensilis–D. abyssinica–D. rotundata* complex, in which species divergence and environmental variation are partly confounded with lineage identity, is very similar to teosinte, where local adaptation to temperature and soil phosphorus contributed to divergence between subspecies despite ongoing gene flow (Aguirre-Liguori et al., 2019). Thus, the yam system requires methods that explicitly account for this confounding, as we have attempted here with the GEAVs. A key question raised by our findings is how candidate loci can be translated into improved prediction of adaptive performance. However, environment-of-origin data are suggested to provide limited benefit for predicting climate adaptation (Li et al., 2025), as the GEAVs here implicitly do. There may be a need to capture allele-specific expression, as this can also impact adaptation (Hu et al., 2022). There are multiple routes to adaptation and implications for yam breeding and this combined framework could be adapted for prioritizing yam wild relatives as donor sources. Yam presents a distinctive opportunity as its combination of a clonally propagated crop with sexually reproducing, partially interfertile wild relatives means that useful alleles identified in *D. praehensilis* or *D. abyssinica* could be introduced by hybridization and then fixed clonally in selected genotypes, as has been proposed for other clonal perennial crops such as grapevine. Realizing this potential will require experimental validation of candidate associations and genomic predictions before they are applied in breeding programs.

GEA methods are increasingly being incorporated into breeding programs to help facilitate the efficient development of cultivars better suited to specific environmental conditions (Forester et al., 2018; Gao et al., 2023; Aitken et al., 2024; Campbell et al., 2025). While GEA provides a useful method for identifying candidate loci and potential parents for breeding, it has limitations, including limited sampling, the resolution of climate and soil data, and population size (Rellstab et al., 2015; Lotterhos & Whitlock, 2015). These limitations were present in our data, sample sizes per group were modest (**Table 1**), and accessions shared collection sites (148 sites for 167 accessions). This meant that there was only the ability to detect loci of large effect (e.g., power simulations in **Figure S7**), so absence of association should not be read as absence of adaptive variation. The very stringent zero-missing-data filter (63,142 of 3,570,940 SNPs), was used because missingness is higher in regions divergent from the *D. rotundata* reference thus this filter probably under-represents variation specific to the wild species, although population structure (Procrustes r = 0.998 for PC1–PC4) and the environment-only RDA fraction (1.79% versus 1.77%) were essentially unchanged with a relaxed, imputed dataset of 1,573,596 SNPs (**Table S3**). The last caveat is that we did not explore structural variants (e.g., presence– absence, copy-number variation, and transposable elements). However, despite these limitations, this study identified loci and accessions that are potentially useful for abiotic stress breeding in Yam.

## Acknowledgements

We would like to thank the reviewers for their constructive comments and suggestions, which greatly improved the quality of the manuscript.

## Author Contributions

Bindu Adhikari - Conceptualization, Formal analysis, Writing, Revising.

Roland Akakpo - Conceptualization, Formal analysis, Writing, Revising.

Anna Halpin-McCormick - Conceptualization, Revising.

Marybel Soto Gomez - Conceptualization, Revising.

Nora Scarcelli - Conceptualization, Revising.

Karine Alix - Conceptualization, Revising.

Yves Vigouroux - Conceptualization, Writing, Revising.

Michael Kantar - Conceptualization, Formal analysis, Writing, Revising.

## Data availability

The resequencing data and SNP calls analyzed here are available from Scarcelli et al. (2019b; https://doi.org/10.5281/zenodo.2540773). Collection-site environmental data, intermediate results and supplementary tables are available at https://github.com/kant0063/Yam_GEA.

## Code availability

Code used for all analyses is archived at https://github.com/kant0063/Yam_GEA

---

**Figure S1.**
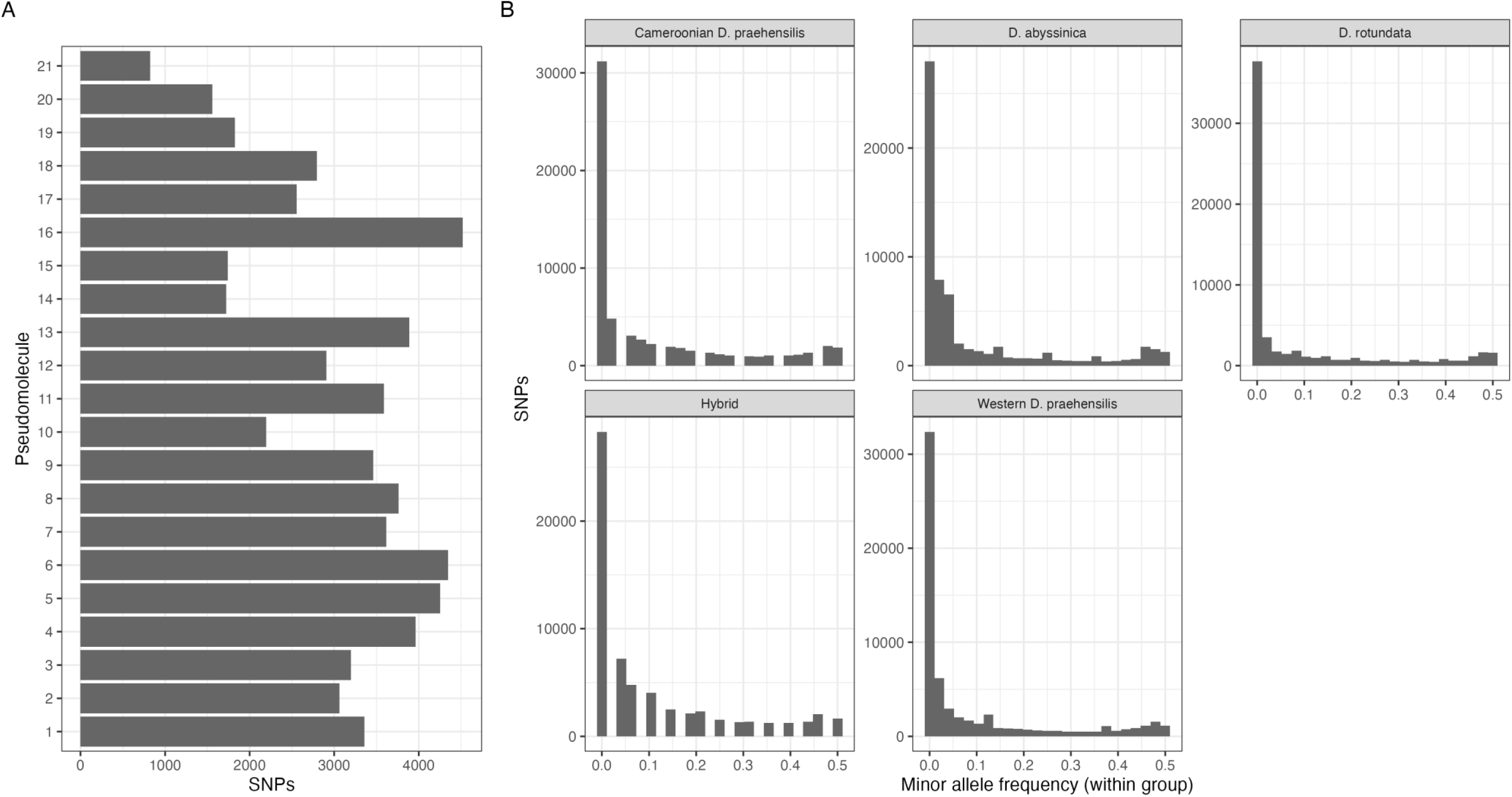
(A) Number of SNPs per pseudomolecule in the zero-missing-data set (63,142 SNPs; 824–4,522 per pseudomolecule). (B) Folded minor allele frequency spectra within each genetic group, including SNPs monomorphic within a group (MAF = 0).

**Figure S2.**
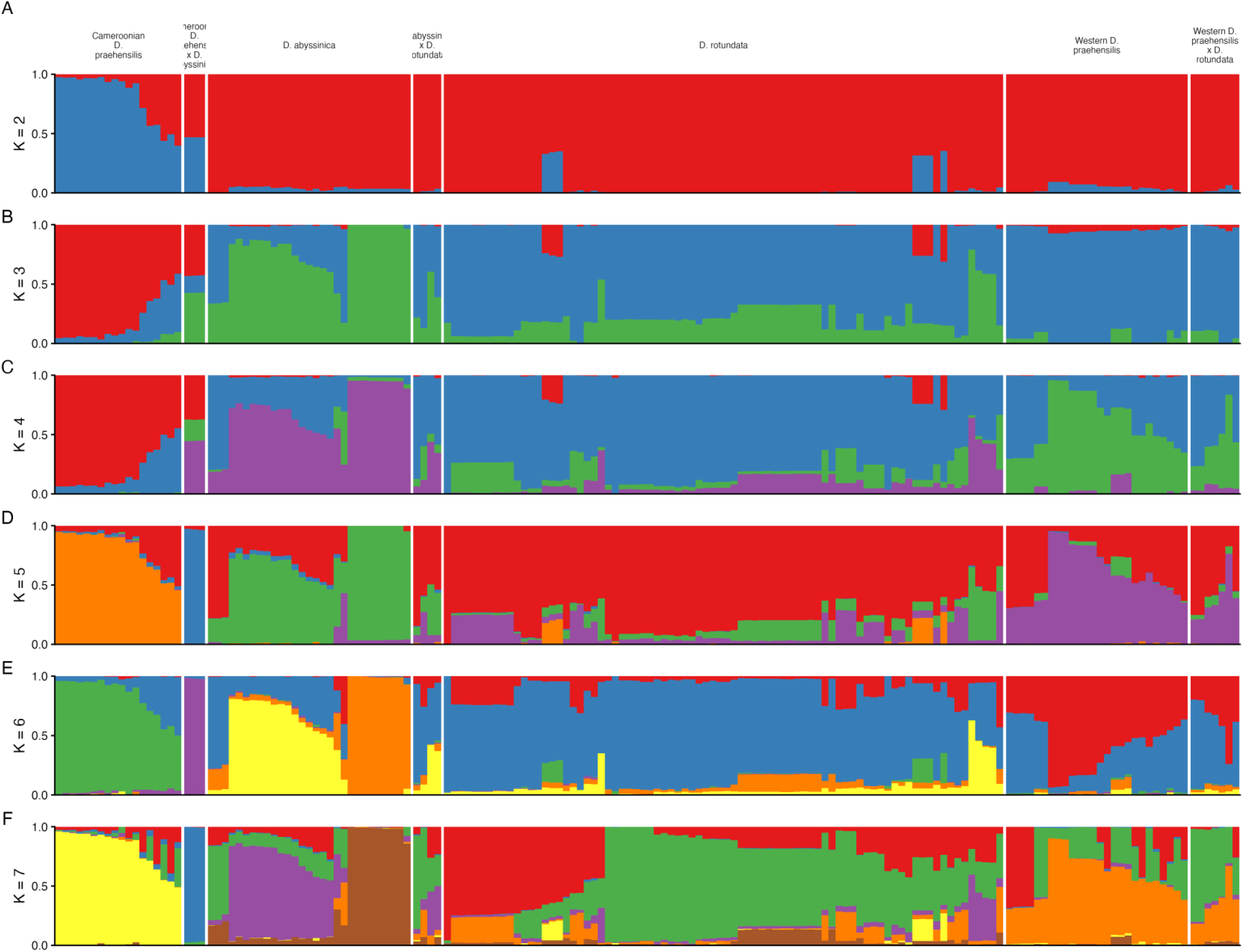
Ancestry proportions estimated with TESS3 for (A) K = 2 to (F) K = 7. Each vertical bar is one accession, colored by its ancestry coefficients; accessions are grouped by genetic group (labels above panel A) and kept in the same order in every panel. Color’s match Figure 1B and Figure S3. Cross-validation supported K = 7, which separates the species and lineages and identifies the hybrids.

**Figure S3.**
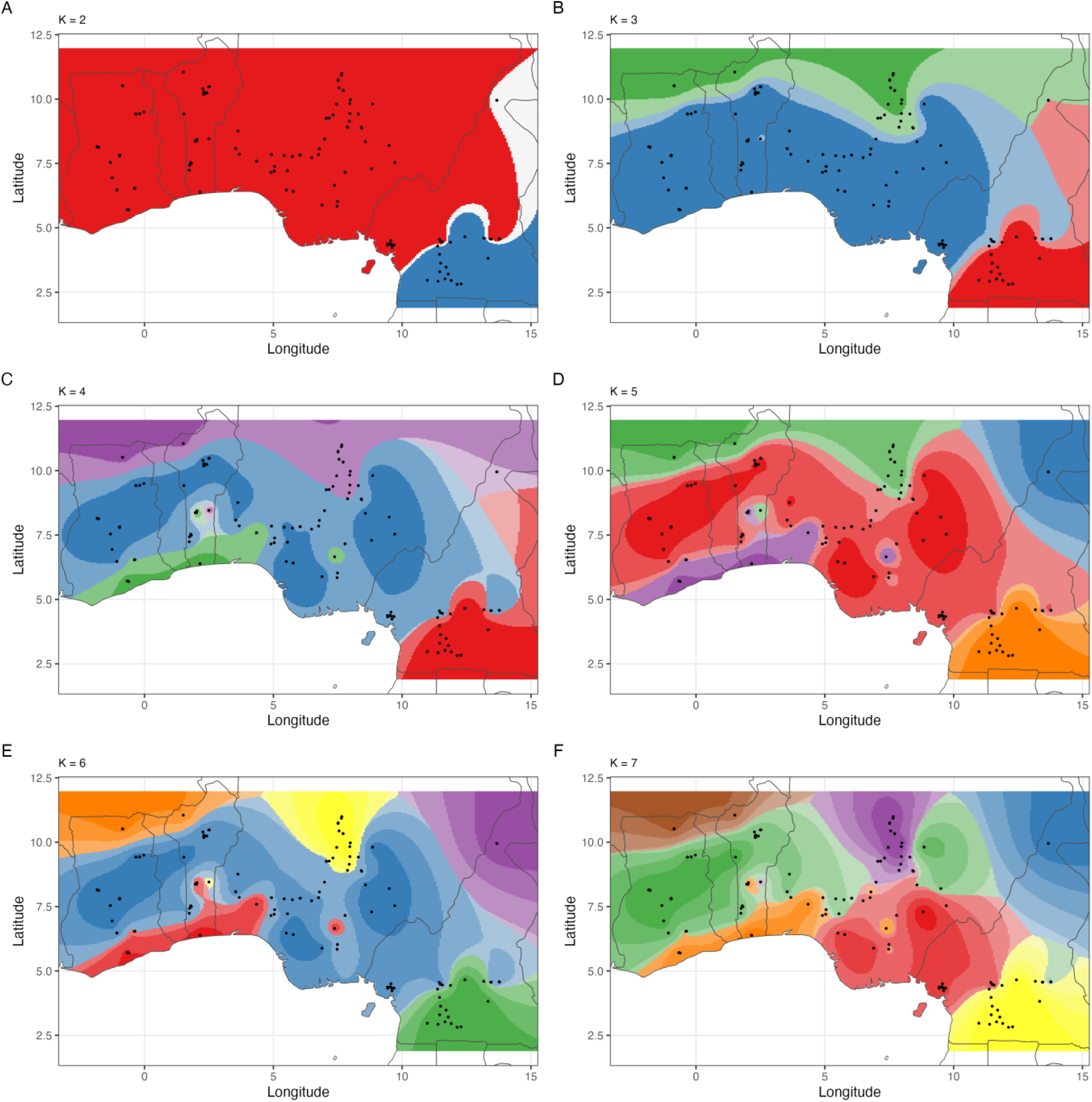
Spatial interpolation of TESS3 ancestry coefficients for (A) K = 2 to (F) K = 7 across the sampled range; black points are collection sites. Color’s match Figure S2, and panel F corresponds to Figure 1C.

**Figure S4.**
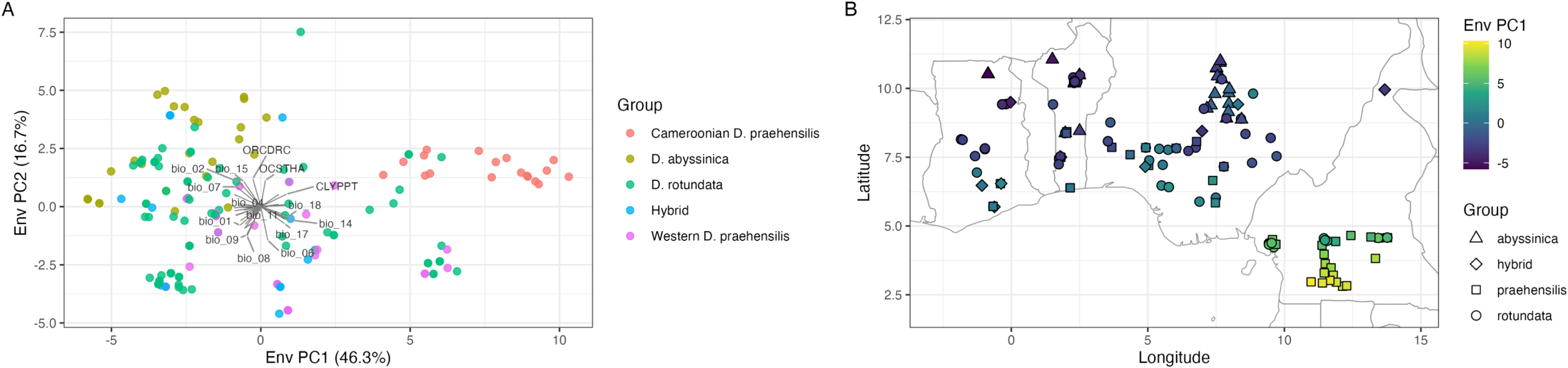
(A) Principal component analysis of the 36 scaled environmental variables at the collection sites of the 167 accessions, colored by genetic group; grey arrows are variable loadings, with the longest labelled (PC1 46.3%, PC2 16.7%). (B) Collection sites colored by their score on environmental PC1, with symbol shape showing species (hybrids separately). Cameroonian *D. praehensilis* sites lie at the high end of PC1 and *D. abyssinica* sites at the low end.

**Figure S5.**
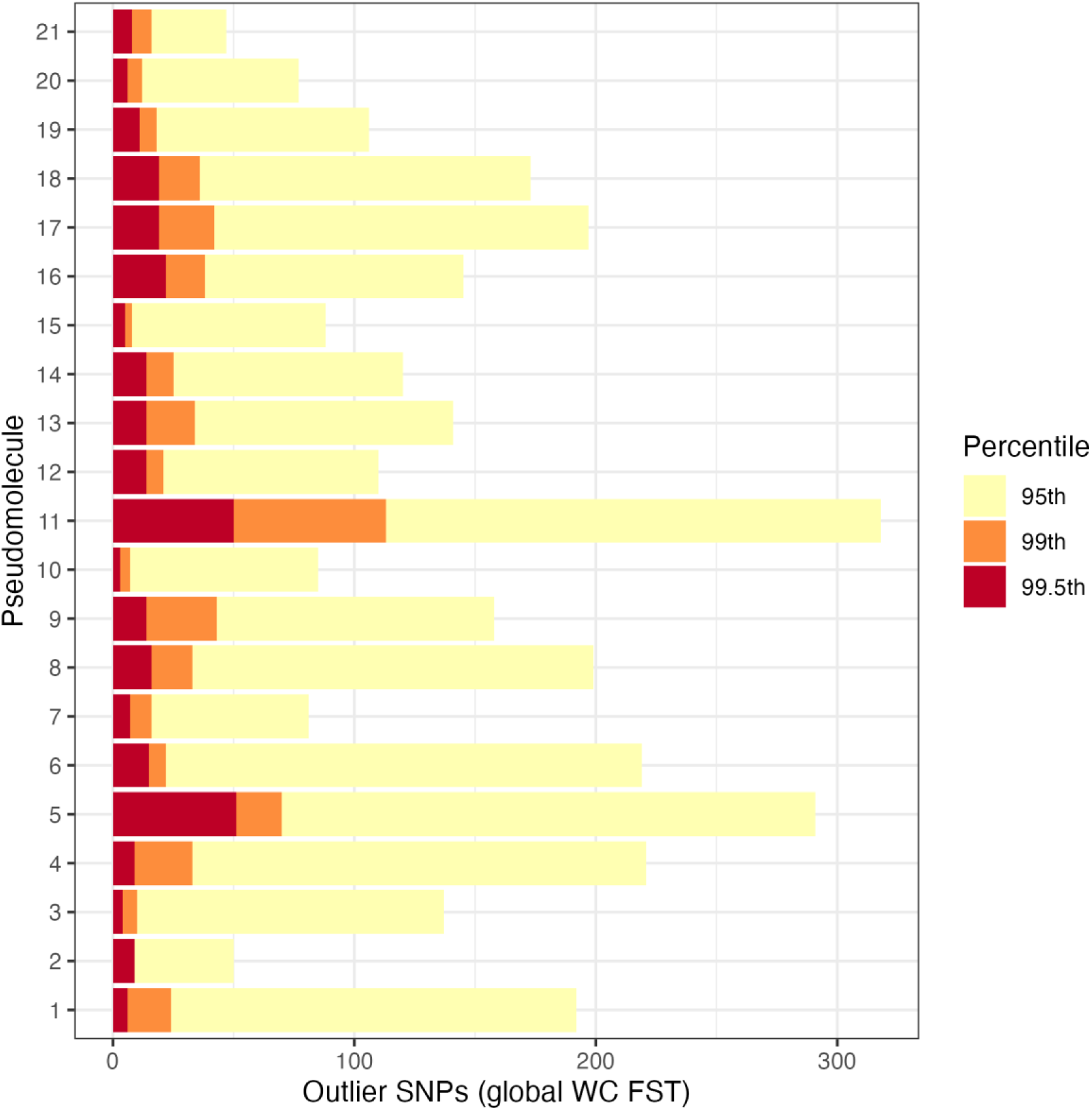
Number of global Weir and Cockerham F_ST_ outlier SNPs per pseudomolecule, by percentile class: 95th– 99th (light yellow), 99th–99.5th (orange) and above the 99.5th percentile (red). Of the 3,155 outliers, 2,525 fell in the 95th–99th, 314 in the 99th–99.5th and 316 above the 99.5th percentile. Outliers were most numerous on BDMI01000011.1 (318) and BDMI01000005.1 (291), which also had the most outliers above the 99.5th percentile (50 and 51), and least numerous on BDMI01000021.1 (47) and BDMI01000002.1 (50). The uneven distribution is not explained by SNP density alone (Figure S1A; BDMI01000016.1 has the most SNPs but 145 outliers) and, like the outliers themselves, mainly reflects differentiation among groups.

**Figure S6.**
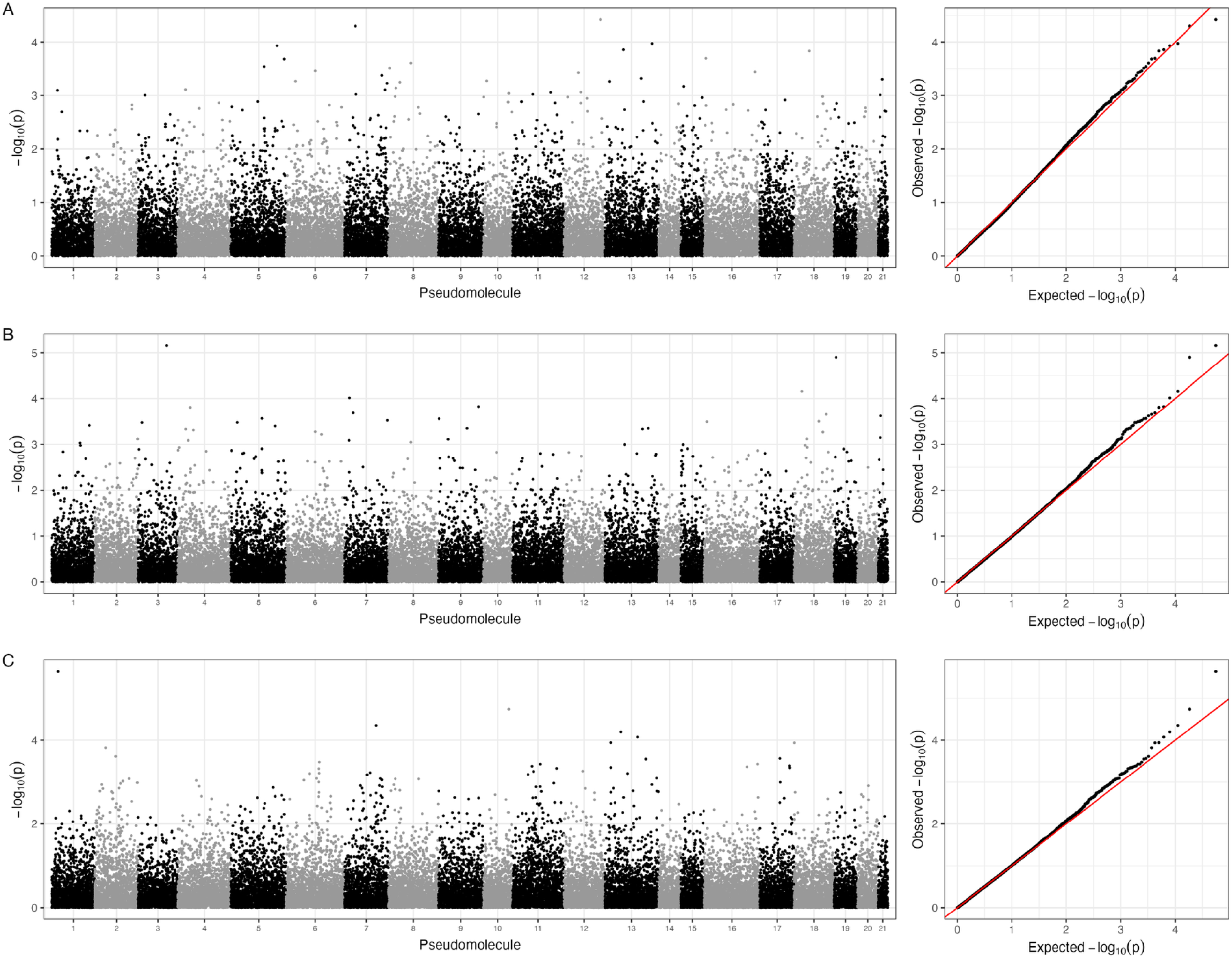
Genome-wide association with the three independent components of the bioclimatic variables (IC1–IC3), using the Q + K mixed model (EMMAX/P3D; kinship and six genetic principal components; 27,785 SNPs). (A) IC1, (B) IC2 and (C) IC3. For each IC, the left panel is a Manhattan plot of −log_10_ p by pseudomolecule (1–21, alternating black and grey) and the right panel a quantile–quantile plot of observed against expected −log_10_ p, with the red line marking the null expectation. Genomic inflation factors were λ = 0.92, 0.93 and 0.97, and no SNP was significant at FDR 0.05 for any IC.

**Figure S7.**
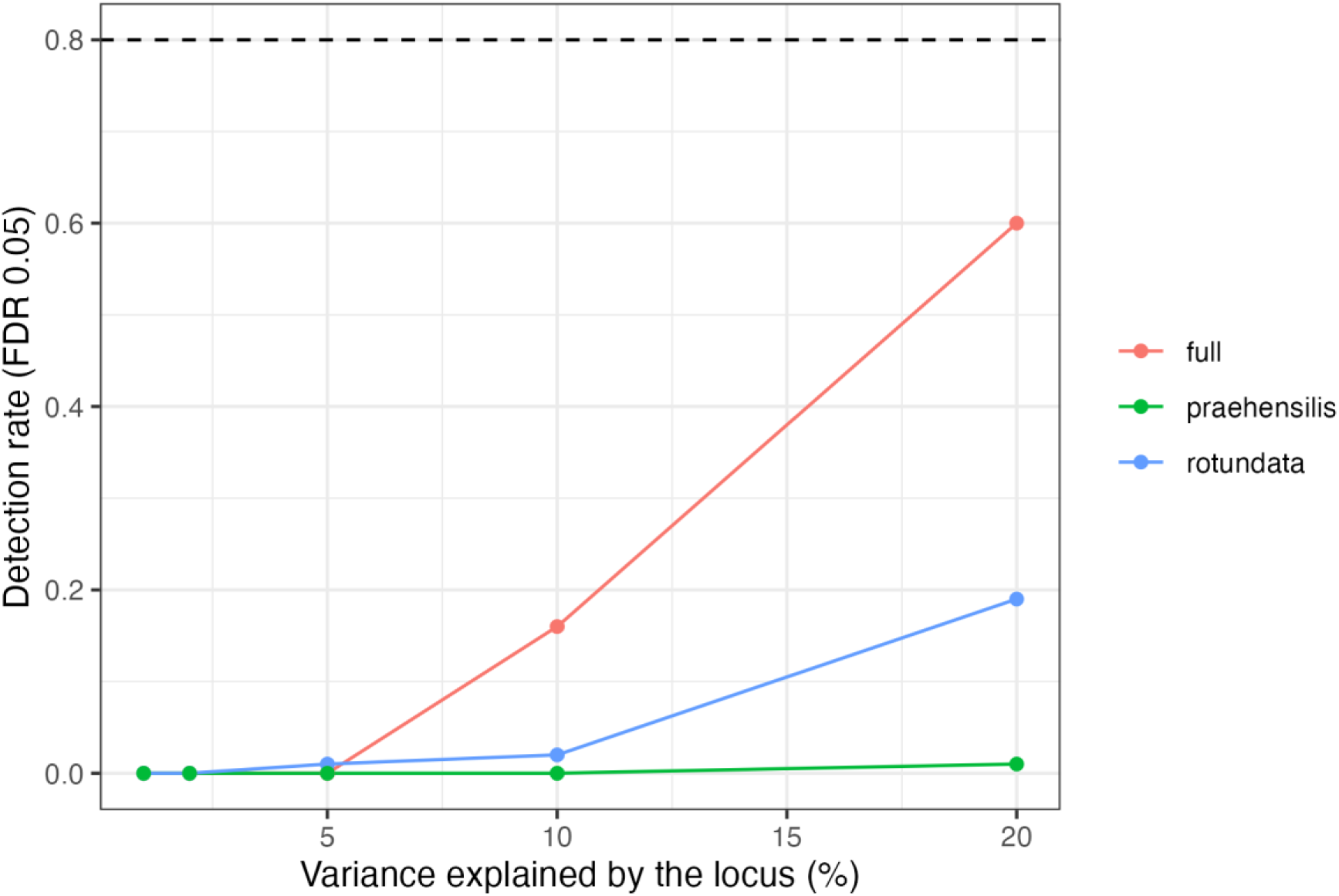
Power to detect a single locus at FDR 0.05 as a function of the proportion of variance it explains (1, 2, 5, 10 and 20%), for the full panel and within *D. rotundata* and *D. praehensilis*, from 100 simulated response variables per scenario on the observed genotypes and kinship, with a polygenic background explaining 30% of the variance. The dashed line marks 80% power.

**Figure S8.**
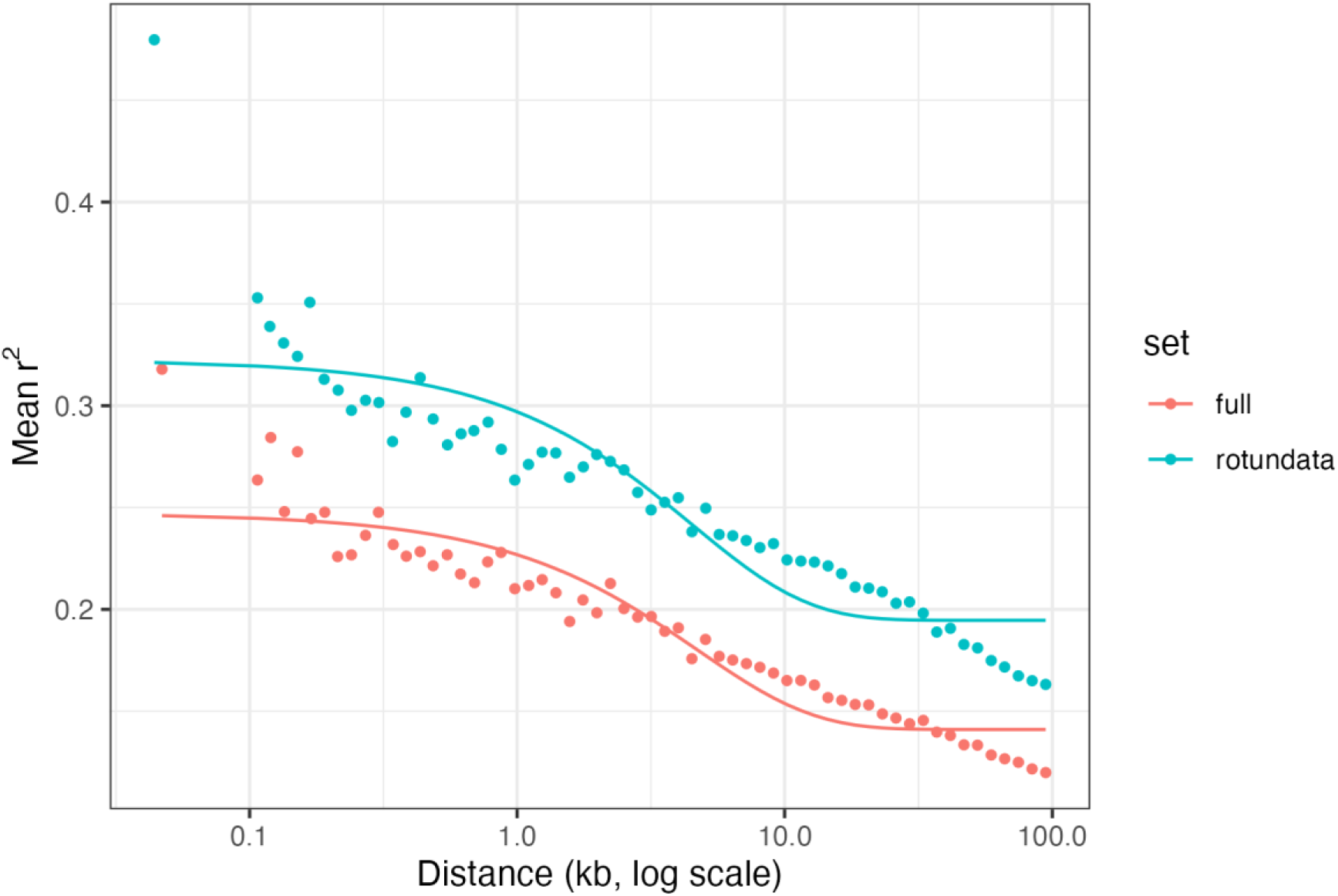
Linkage disequilibrium decay in the full panel (red) and within *D. rotundata* (teal), estimated from the relaxed, imputed SNP set (1.19 million SNP pairs in the full panel and 0.78 million within *D. rotundata*). Points are mean r² in log-spaced distance bins for SNP pairs up to 100 kb apart, and lines are the fitted model r² = c + a·exp(−bd). The half-decay distance was 3,275 bp in the full panel and 3,123 bp within *D. rotundata*; fitted r² fell to 0.2 at 2.8 kb and 14.3 kb, respectively, with fitted background r² of 0.14 and 0.19. Beyond about 30 kb, observed r² continued to decline below the fitted background, consistent with long-range LD generated by population structure.

**Figure S9.**
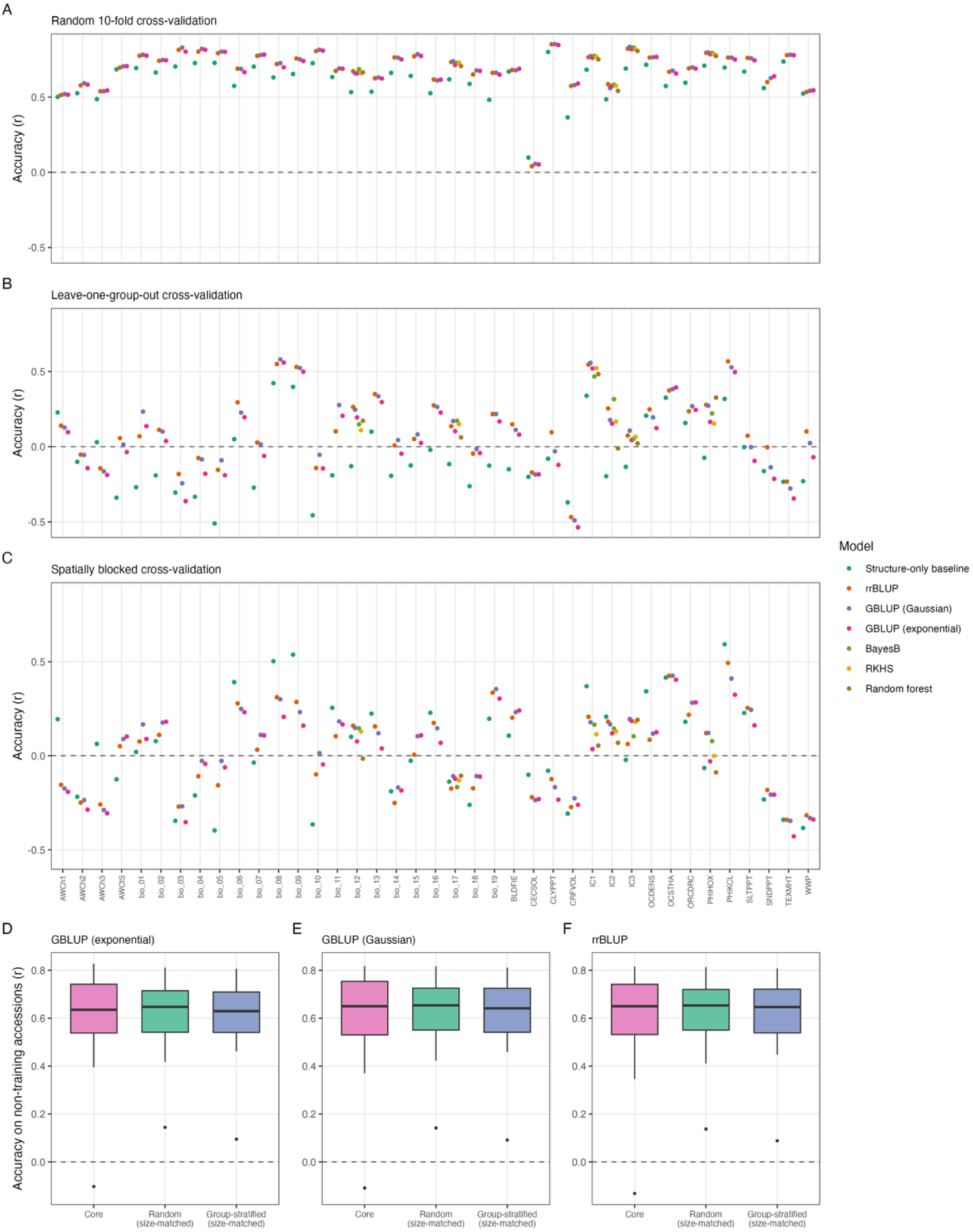
Accuracy of environmental genomic prediction (Pearson’s r between out-of-fold predictions and observed values) for the 36 environmental variables and three ICs under (A) random 10-fold (five repetitions), (B) leave-one-group-out and (C) spatially blocked (six k-means blocks of coordinates) cross-validation using all 167 accessions. Colours show models: the structure-only baseline (regression on six genetic principal components), rrBLUP and GBLUP with Gaussian and exponential kernels for all variables, and BayesB, RKHS and random forest (RF) for IC1–IC3, BIO12, BIO17 and PHIHOX only. The dashed line marks r = 0. Mean accuracy of the genomic models across variables was 0.69–0.73 under random cross-validation (baseline 0.61), 0.05–0.23 under leave-one-group-out (baseline −0.08) and 0.01–0.08 under spatial blocks (baseline 0.04); means for BayesB, RKHS and RF are over six variables only. (D–F) Accuracy on non-training accessions when training on the 50-accession core selected from genotypes (Core) or on 50 random (Random) or 50 genetic-group-stratified (Group-stratified) training sets of the same size, for (D) GBLUP with an exponential kernel, (E) GBLUP with a Gaussian kernel and (F) rrBLUP. Boxes summarize the 39 response variables (median, interquartile range, whiskers at 1.5 × the interquartile range); for the random and stratified designs, each value is the mean over the 50 training sets. The dashed line marks r = 0. Results for the original core, which contained no Cameroonian *D. praehensilis* and performed much worse, are in Table S15.

**Figure S10.**
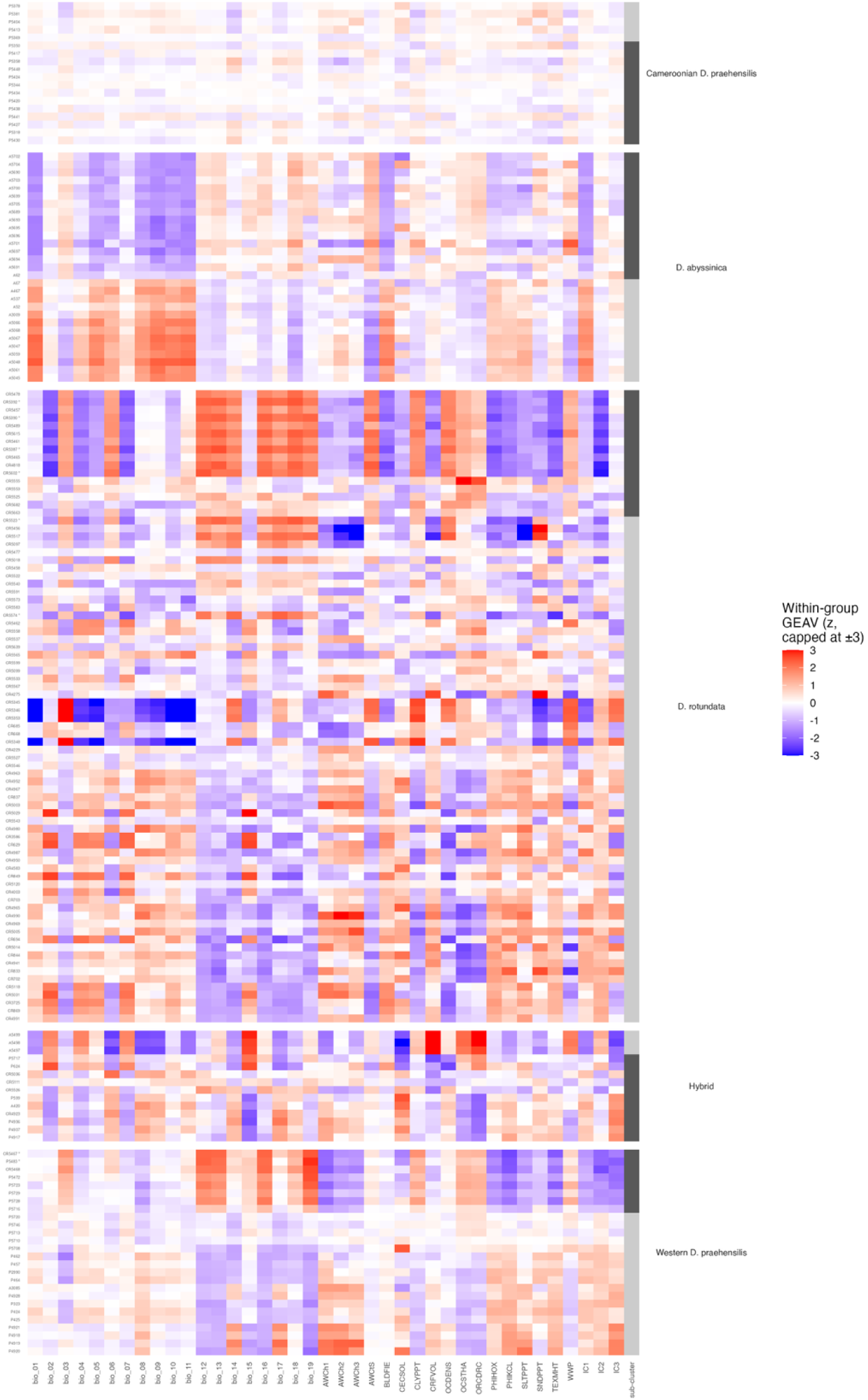
Within-group genomic estimated adaptive values (GEAVs) for the 36 environmental variables and three ICs (columns) and 167 accessions (rows), from out-of-fold predictions expressed as deviations from each accession’s genetic-group mean and standardized (z-scores; color scale capped at ±3). Accessions (rows, labelled) are grouped by genetic group (labels at right), the bar at right marks two sub-clusters within each group (dark and light grey), defined by k-means clustering (k = 2) of the first two principal components of within-group genotypes; rows are ordered within groups by the first of these components, and asterisks mark the candidate parents named in the text. Red indicates values above and blue values below the group mean. Within-group GEAVs vary least in Cameroonian *D. praehensilis*, whereas the two *D. abyssinica* sub-clusters have opposite predicted values for the temperature variables (bio_01–bio_11).

**Figure S11.**
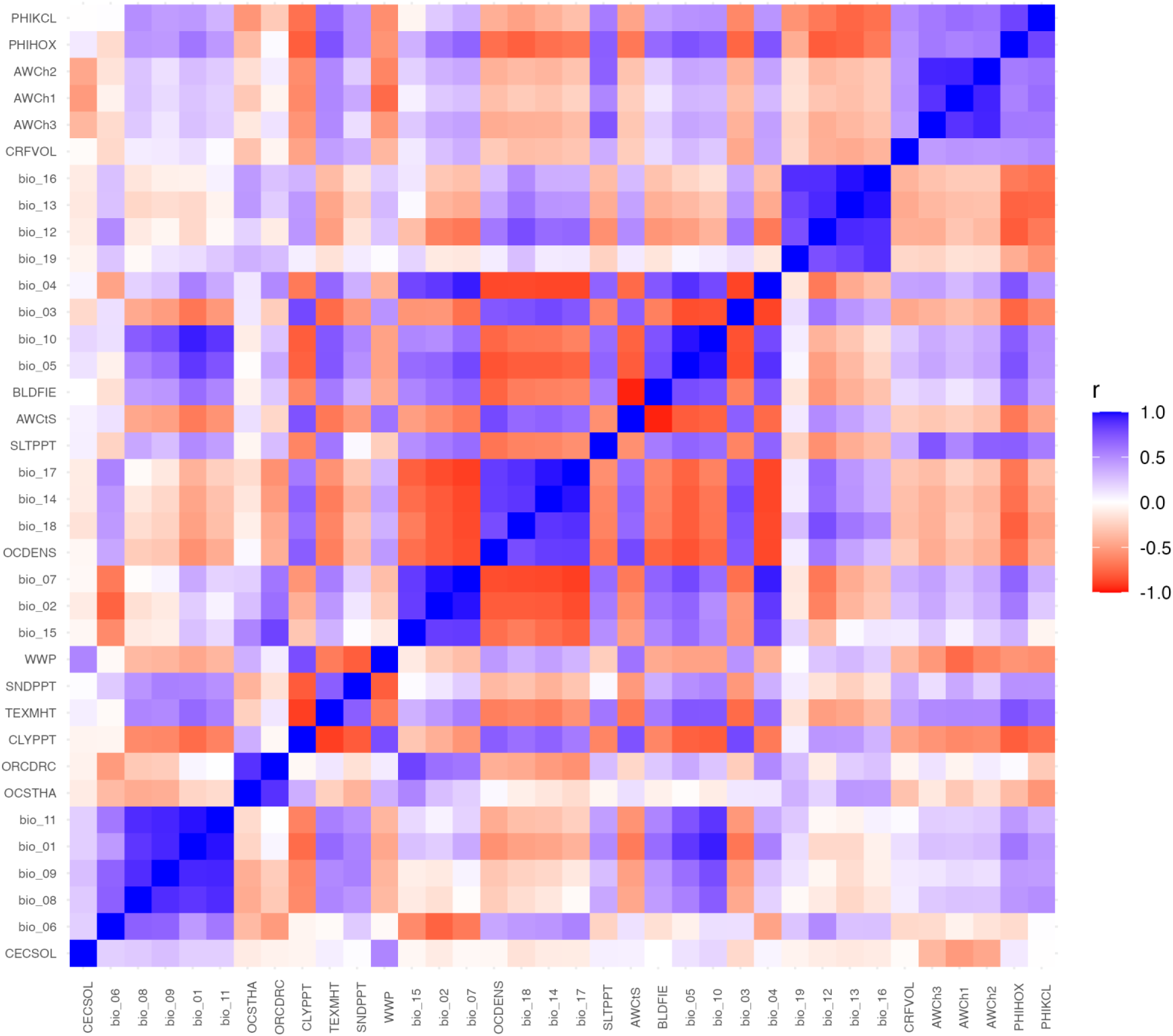
Pearson correlations among the 36 environmental variables at the collection sites: 19 bioclimatic variables (bio_01–bio_19) and 17 soil variables (AWCh1–AWCh3, available soil water capacity at three matric potentials; AWCtS, saturated water content; BLDFIE, bulk density; CECSOL, cation exchange capacity; CLYPPT, SLTPPT and SNDPPT, clay, silt and sand content; CRFVOL, coarse fragment volume; OCDENS, organic carbon density; OCSTHA, organic carbon stock; ORCDRC, organic carbon content; PHIHOX and PHIKCL, pH in water and in KCl; TEXMHT, texture class; WWP, available water capacity until wilting point). Variables are ordered so that correlated variables are adjacent; blue indicates positive and red negative correlations. Annual and wet-season precipitation (bio_12, bio_13, bio_16, bio_19) form one positively correlated block, and dry-season precipitation (bio_14, bio_17, bio_18) and OCDENS another, which is negatively correlated with temperature range and seasonality (bio_02, bio_04, bio_07) and with precipitation seasonality (bio_15). Clay content (CLYPPT) is negatively correlated with sand content and texture (SNDPPT, TEXMHT).

**Figure S12.**
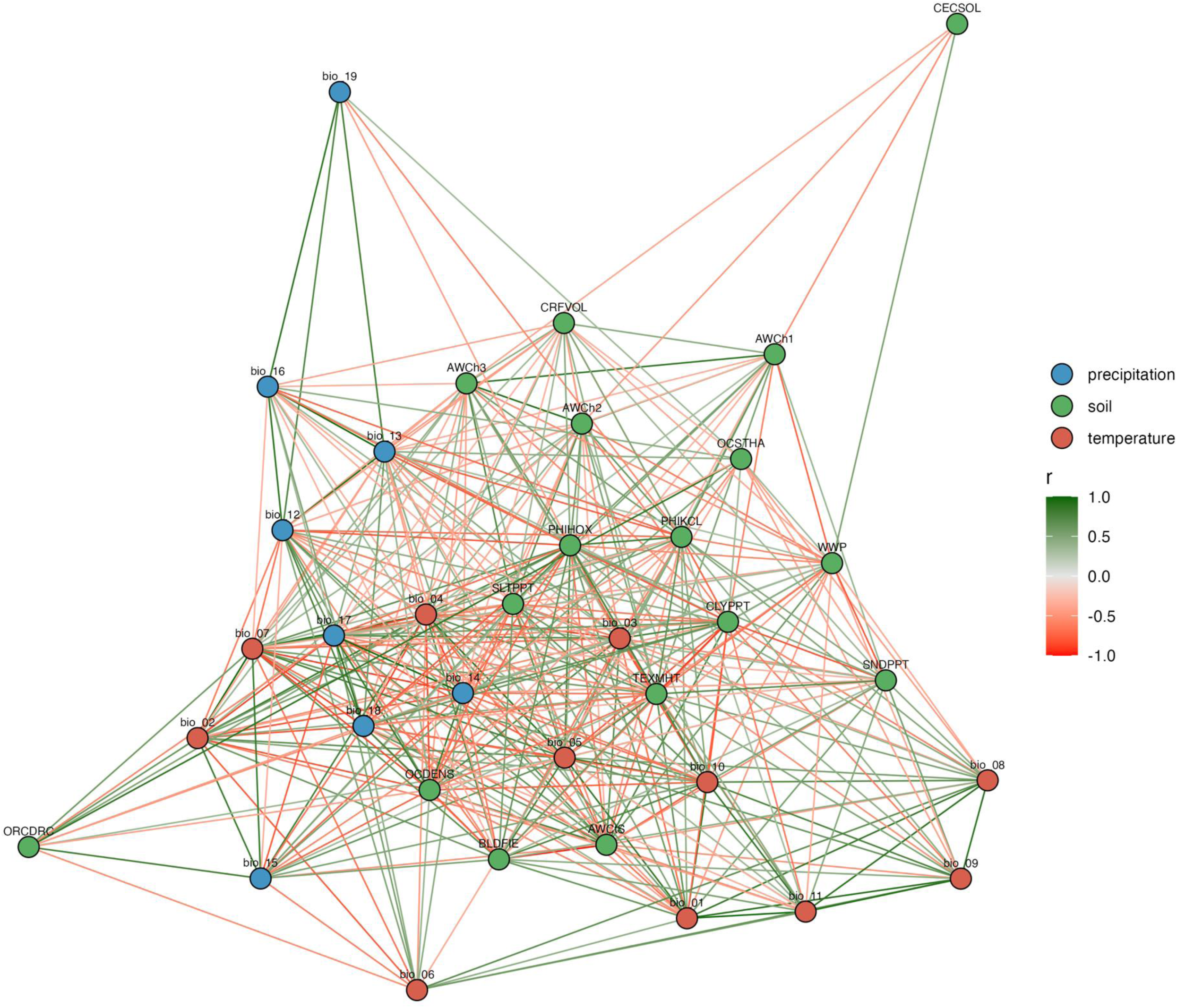
Correlation network of the 36 environmental variables. Nodes are variables colored by type (blue, precipitation; red, temperature; green, soil); edges connect pairs with |r| ≥ 0.3 and are green for positive and red for negative correlations, with intensity proportional to |r|. Node positions come from a force-directed layout in which strongly correlated variables lie close together. Annual and wet-season precipitation (bio_12, bio_13, bio_16, bio_19) lie together at upper left, dry-season precipitation (bio_14, bio_17, bio_18) lies with organic carbon density (OCDENS), mean and maximum temperatures (bio_01, bio_05, bio_08–bio_11) lie at lower right, and soil texture and pH variables occupy the center. CECSOL and ORCDRC have the fewest connections.

**Figure S13.**
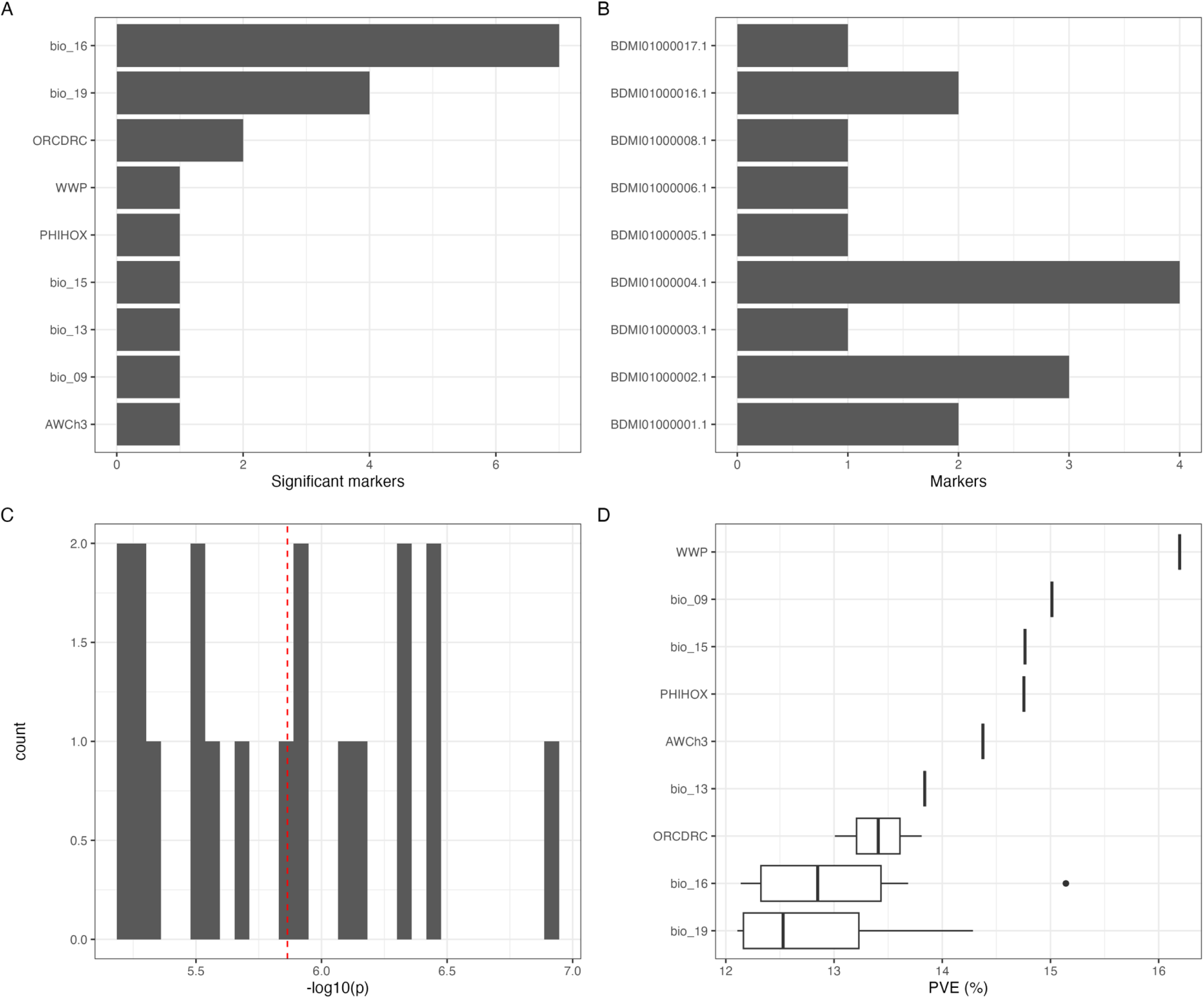
Summary of the 19 full-panel marker–environment associations at 16 markers (FDR 0.05). (A) Number of significant markers per environmental variable; associations were found for nine variables, most often precipitation of the wettest quarter (bio_16; 7 markers) and of the coldest quarter (bio_19; 4). (B) Number of significant markers per pseudomolecule (nine pseudomolecules). (C) Distribution of association strength (−log_10_ p) across the 19 associations; the red dashed line is the median. (D) Proportion of residual variance explained (PVE, %) by significant markers for each variable (12.1–16.2%).

**Figure S14.**
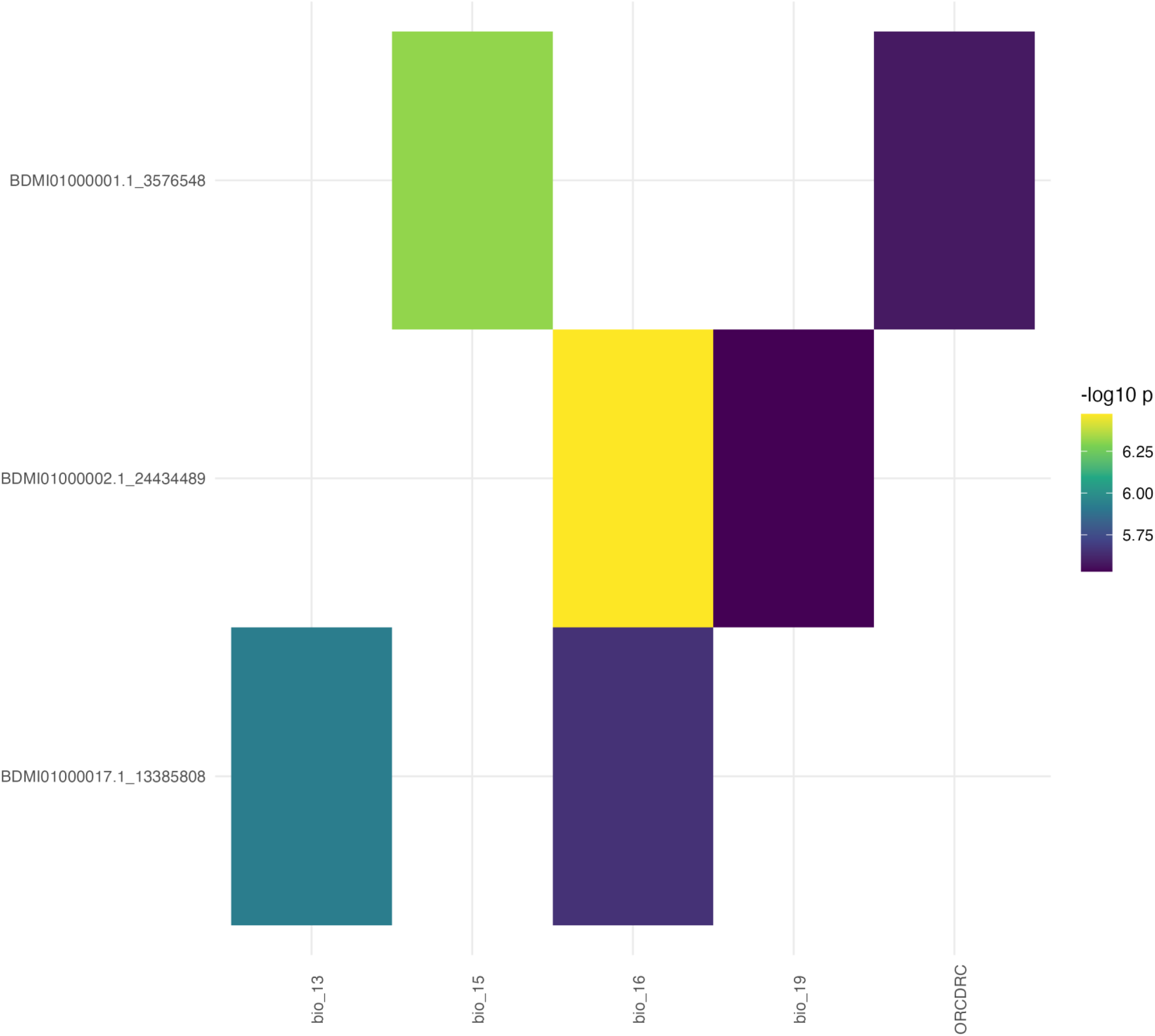
Association strength (−log_10_ p) for the three full-panel markers associated with more than one environmental variable; blank cells are non-significant marker–variable combinations. BDMI01000001.1_3576548 was associated with precipitation seasonality (bio_15) and soil organic carbon content (ORCDRC), BDMI01000002.1_24434489 with precipitation of the wettest and coldest quarters (bio_16, bio_19), and BDMI01000017.1_13385808 with precipitation of the wettest month and wettest quarter (bio_13, bio_16). Because these variables are correlated (Figures S11–S12), multiple associations may reflect a single gradient rather than pleiotropy.

**Figure S15.**
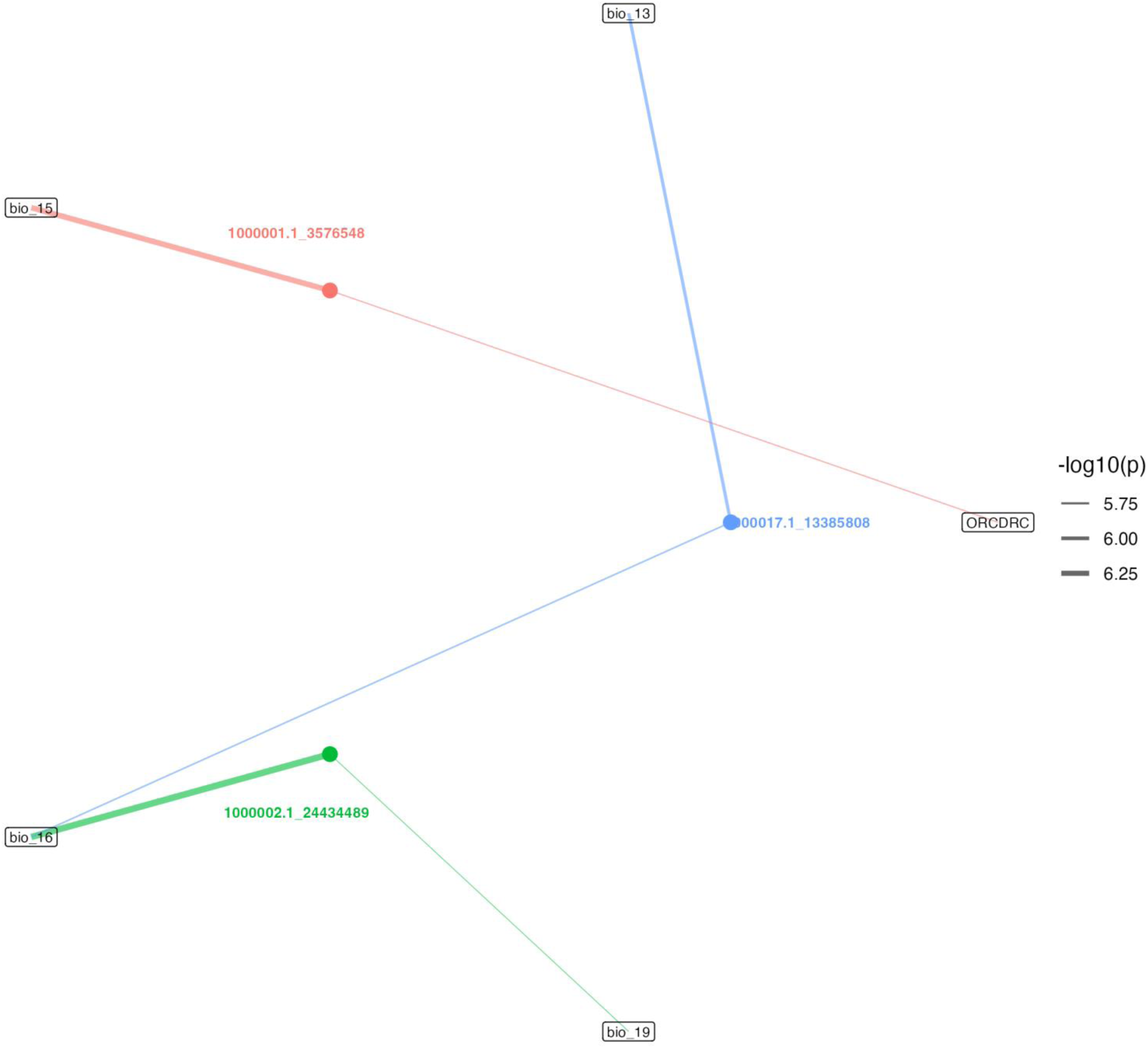
Network of the associations shown in Figure S14. Points are the three markers associated with more than one variable (red, BDMI01000001.1_3576548; green, BDMI01000002.1_24434489; blue, BDMI01000017.1_13385808) and labels are the associated variables; line width is proportional to −log_10_ p.

**Figure S16.**
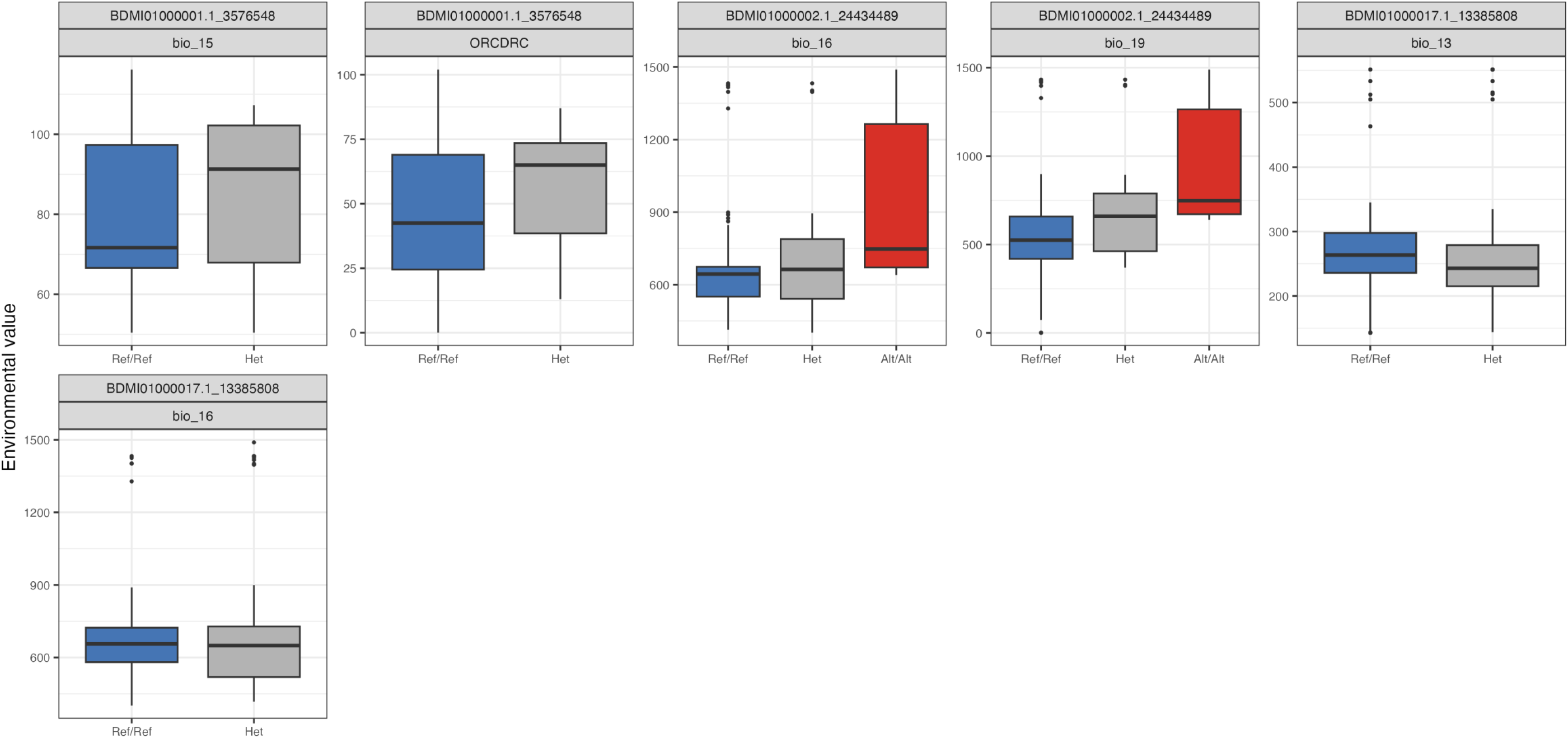
Environmental values at collection sites by genotype for the six associations of the three full-panel markers associated with more than one environmental variable (marker, upper strip; variable, lower strip): BDMI01000001.1_3576548 with precipitation seasonality (bio_15) and soil organic carbon content (ORCDRC), BDMI01000002.1_24434489 with precipitation of the wettest and coldest quarters (bio_16, bio_19), and BDMI01000017.1_13385808 with precipitation of the wettest month and wettest quarter (bio_13, bio_16). Genotypes are Ref/Ref (blue), heterozygous (grey) and Alt/Alt (red); no homozygous alternate genotypes were observed at BDMI01000001.1_3576548 or BDMI01000017.1_13385808. Values are in original units (bio_15, coefficient of variation, %; ORCDRC, g kg^−1^; precipitation, mm). Boxplots show the median, interquartile range and whiskers at 1.5 × the interquartile range. Allele states also covary with genetic group, so these differences are consistent with, but do not demonstrate, local adaptation.

**Figure S17.**
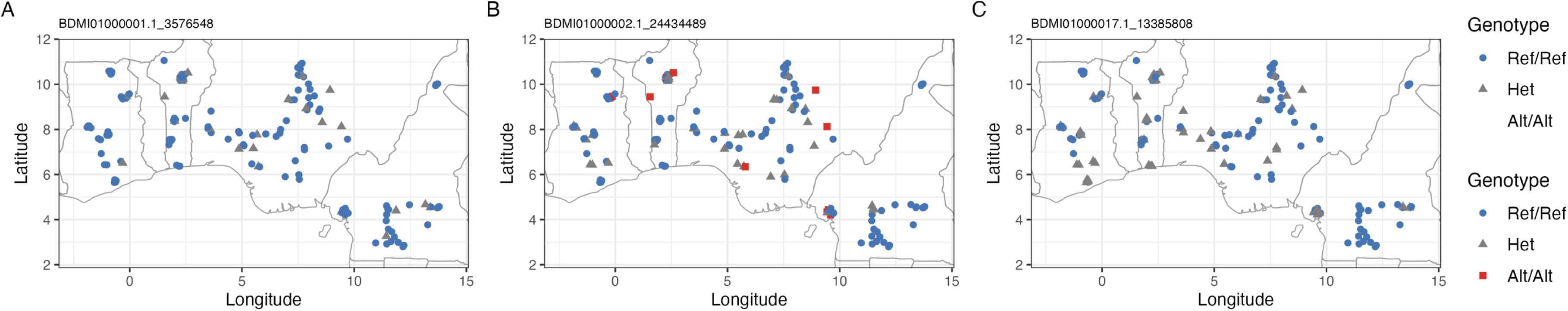
Geographic distribution of genotypes at the three full-panel markers associated with more than one environmental variable ((A) BDMI01000001.1_3576548, (B) BDMI01000002.1_24434489 and (C) BDMI01000017.1_13385808): Ref/Ref (blue circles), heterozygous (grey triangles) and Alt/Alt (red squares). Homozygous alternate genotypes occurred only at BDMI01000002.1_24434489, mostly in Benin and Nigeria. Alternative alleles were less common at Cameroonian sites, so genotypes also track genetic group, and these patterns are consistent with, but do not by themselves demonstrate, local adaptation.

**Figure S18.**
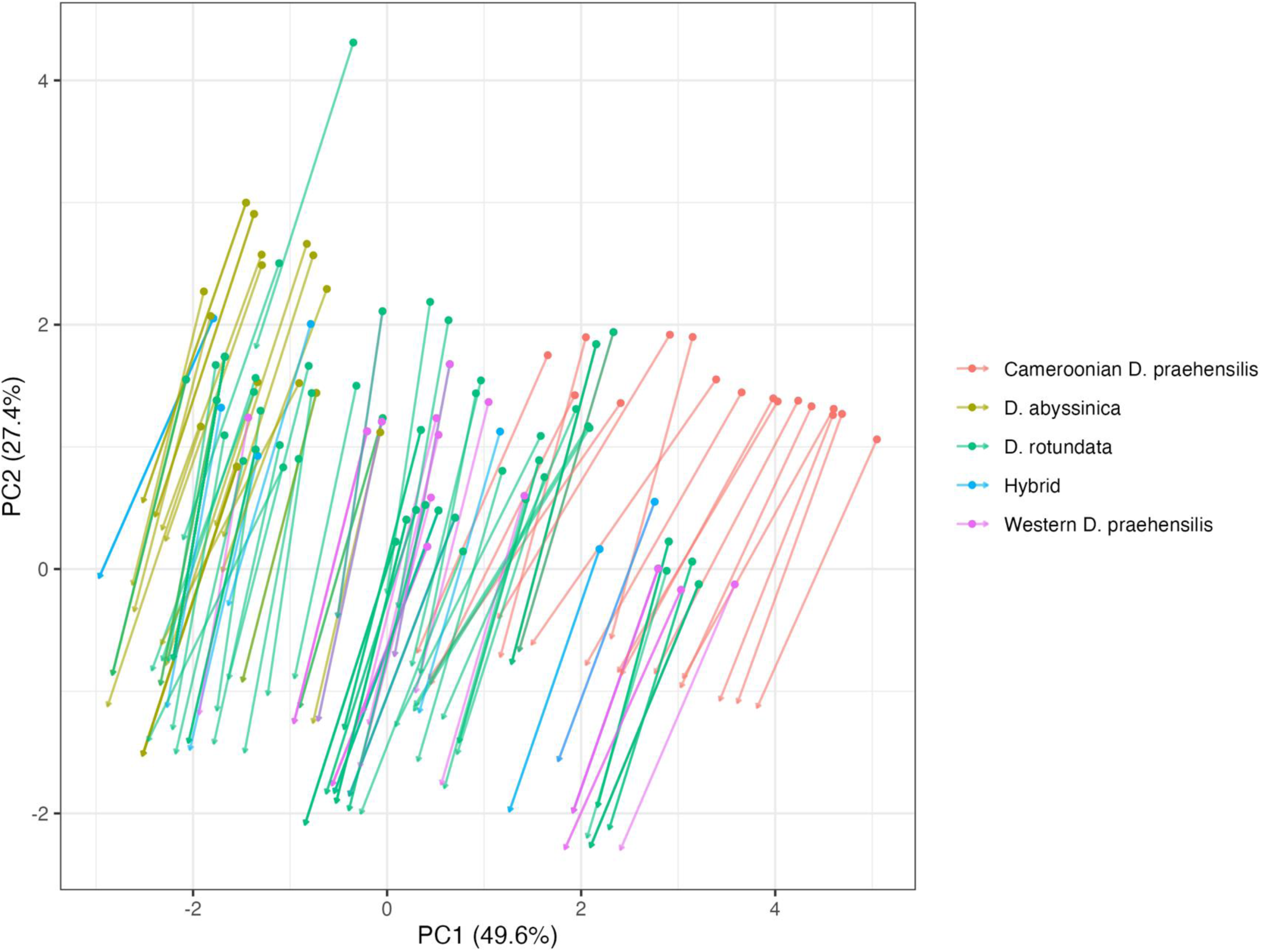
Shift in bioclimate at the sampling sites between current conditions (WorldClim 2.1, 1970–2000; points) and SSP5-8.5, 2081–2100 (mean of six CMIP6 models; arrow heads), shown on the first two principal components of the seven bioclimatic variables used for genomic offsets (PC1 49.6%, PC2 27.4%). Points and arrows are colored by genetic group. All sites shift in a similar direction, mainly toward lower PC2 scores.

**Table S1.** Environmental variables: definitions, units and sources, and whether each was retained after VIF pruning (16 variables) and used in the genomic offset analysis (seven bioclimatic variables).

**Table S2.** Effect of the zero-missing-data filter by genetic group: number of accessions, percentage of SNPs with at least one missing call, mean missing rate, total SNPs and SNPs retained by the zero-missing filter.

**Table S3.** Comparison of key results between the zero-missing-data set (63,142 SNPs) and the relaxed dataset (≤ 20% missing, MAF ≥ 0.05, imputed with Beagle; 1,573,596 SNPs): Procrustes correlation of genetic PCs 1–4, the environment-only pRDA fraction, the number of full-panel EGWAS associations and the number of associated 10-kb windows shared across the full panel and both within-species sets.

**Table S4.** Isolation by distance and by environment: MMRR coefficients and partial Mantel tests for the full panel and within *D. rotundata*, *D. praehensilis* and *D. abyssinica*.

**Table S5.** Loadings of the 19 bioclimatic variables on the three independent components (IC1–IC3) used as summary climate variables.

**Table S6.** WZA windows in the top 0.1% within *D. rotundata* and *D. praehensilis*, with overlap with EGWAS candidates.

**Table S7.** Genomic inflation factor (λ), null-model heritability, number of SNPs tested and number of significant associations (FDR 0.05) for every environmental variable and IC, for the full panel and within *D. rotundata* and *D. praehensilis*; “boundary” marks fits in which the variance component reached its bound.

**Table S8.** LFMM significant SNPs (FDR 0.05) by environmental variable for the full panel (K = 7) and within *D. rotundata* (K = 3) and *D. praehensilis* (K = 2).

**Table S9.** BSLMM posterior estimates of PVE, PGE and number of sparse-effect loci (n) for IC1–IC3 and selected variables. Estimates for each of the three MCMC chains are in Table S10.

**Table S10.** BSLMM estimates of PVE, PGE and number of sparse-effect loci (n) for each of the three MCMC chains, for the variables and sets in Table S9.

**Table S11.** Associated markers and their priority score for candidate-gene search: 2 if detected within a species, plus 1 each for a top WZA window, association with more than one variable and detection by LFMM (possible range 0– 5; observed maximum 3), with the evidence behind each score.

**Table S12.** Gene and mRNA features in the TDr96 annotation within 10 kb of markers scoring at least 2 in Table S11, with the associated variables, analysis set and distance from the marker (0, marker within the gene). For the two multi-variable full-panel markers annotated in detail, the position class within the gene and the best Araport11 (*Arabidopsis thaliana*) match from reciprocal BLASTP are also given.

**Table S13.** Per-accession genomic offsets (mean and among-model standard deviation across six CMIP6 models, SSP5-8.5, 2081–2100) for the full panel and within *D. rotundata* and *D. praehensilis*, with Köppen–Geiger classes for 1991–2020 and 2071–2099 at each collection site.

**Table S14.** Accessions in the two 50-accession cores compared as training sets: the core selected from genotypes with Core Hunter and used as the training set (23 *D. rotundata*, 8 western and 7 Cameroonian *D. praehensilis*, 3 *D. abyssinica* and 9 hybrids) and the original core (18 *D. rotundata*, 13 western *D. praehensilis*, 12 *D. abyssinica* and 7 hybrids; no Cameroonian *D. praehensilis*), with genetic group and collection coordinates. Eleven accessions are in both cores.

**Table S15.** Prediction accuracy of the original and genotype-only 50-accession cores used as training sets.

**Table S16.** Within-group genomic estimated adaptive values (GEAVs; out-of-fold predictions expressed as deviations from the genetic-group mean) for 167 accessions, 36 environmental variables and three ICs (Table S17 lists the top-ranked candidate parents).

**Table S17.** Top-ranked candidate parents within each genetic group for low soil pH (PHIHOX) and high annual precipitation (BIO12), with their within-group GEAVs (Table S16).

**Table S18.** Environmental Genome Wide Association Significant Markers with allelic effect (β), standard error, allele frequency, proportion of variance explained (PVE), q-value and genomic inflation factor (λ), for the full panel and within *D. rotundata* and *D. praehensilis*.

**Table S19.** GenoBridge best model and hold-out accuracy (34 accessions) for all 39 variables, compared with the best model and the structure-only baseline under random 10-fold cross-validation.

## Abbreviations

GEA: Genome-Environment Association
EGWAS: Environmental Genome-Wide Association Study
GWAS: Genome-Wide Association Study
FIGS: Focused Identification of Germplasm Strategy
RDA: Redundancy Analysis
WZA: Weighted-Z Analysis
F_ST_: Fixation index (measure of population differentiation)
FROH: Inbreeding coefficient based on runs of homozygosity
ROH: Runs of Homozygosity
LD: Linkage Disequilibrium
PCA: Principal Component Analysis
IC/ICs: Independent Component(s)
ICA: Independent Component Analysis
FDR: False Discovery Rate
LFMM: Latent Factor Mixed Model
EGS: Environmental Genomic Selection
GEAVs: Genomic Estimated Adaptive Values
GEBVs: Genomic Estimated Breeding Values
SNP / SNPs: Single Nucleotide Polymorphism(s)
VCF: Variant Call Format (file)
BLAST: Basic Local Alignment Search Tool
SSP: Shared Socioeconomic Pathway (climate scenario)
CMIP6: Coupled Model Intercomparison Project Phase 6
BIO1–BIO19: WorldClim bioclimatic variable codes
CLYPPT: Clay percentage (soil variable, ISRIC)
CRFVOL: Coarse fragment volume (soil variable, ISRIC)
OCDENS: Organic carbon density (soil variable, ISRIC)
ORCDRC: Organic carbon content (soil variable, ISRIC)
PHIHOX: Soil pH (ISRIC)
TEXMHT: Soil texture (ISRIC)
WWP / AWCh1–3: Available soil water capacity until wilting point / available soil water capacity at three matric potentials (soil variables, ISRIC)
AWCtS: Saturated water content (soil variable, ISRIC)
BLDFIE: Bulk density (soil variable, ISRIC)
CECSOL: Cation exchange capacity (soil variable, ISRIC)
OCSTHA: Organic carbon stock (soil variable, ISRIC)
PHIKCL: Soil pH in KCl (ISRIC)
SLTPPT / SNDPPT: Silt / sand content (soil variables, ISRIC)
K: Number of genetic clusters (population structure analysis)
BSLMM: Bayesian Sparse Linear Mixed Model
CV: Cross-Validation
EMMAX: Efficient Mixed-Model Association eXpedited
ExDet: Extrapolation Detection
IBD / IBE: Isolation by Distance / Isolation by Environment
MMRR: Multiple Matrix Regression with Randomization
pRDA: partial Redundancy Analysis
PGE / PVE: Proportion of genetic variance explained by sparse effects / Proportion of variance explained
VIF: Variance Inflation Factor

